# Schwann cells modulate nociception in neurofibromatosis 1

**DOI:** 10.1101/2023.03.18.533004

**Authors:** Namrata G.R. Raut, Laura A. Maile, Leila M. Oswalt, Irati Mitxelena, Aaditya Adlakha, Kourtney L. Sprague, Ashley R. Rupert, Lane Bokros, Megan C. Hofmann, Jennifer Patritti-Cram, Tilat A. Rizvi, Luis F. Queme, Kwangmin Choi, Nancy Ratner, Michael P. Jankowski

**Affiliations:** Department of Anesthesia, Division of Pain Management, Cincinnati Children’s Hospital Medical Center, Cincinnati, OH 45229; Department of Pediatrics, University of Cincinnati College of Medicine, Cincinnati, Ohio 45229; Pediatric Pain Research Center, Cincinnati Children’s Hospital Medical Center, Cincinnati, OH 45229; Graduate Program in Neuroscience, University of Cincinnati College of Medicine, Cincinnati, Ohio 45229; Division of Cancer Biology and Experimental Hematology, Cincinnati Children’s Hospital Medical Center, Cincinnati, OH 45229

## Abstract

Pain of unknown etiology is frequent in individuals with the tumor predisposition syndrome Neurofibromatosis 1 (NF1), even when tumors are absent. Schwann cells (SC) were recently shown to play roles in nociceptive processing, and we find that chemogenetic activation of SCs is sufficient to induce afferent and behavioral mechanical hypersensitivity in mice. In mouse models, animals show afferent and behavioral hypersensitivity when SC, but not neurons, lack *Nf1*. Importantly, hypersensitivity corresponds with SC-specific upregulation of mRNA encoding glial cell line derived neurotrophic factor (GDNF), independent of the presence of tumors. Neuropathic pain-like behaviors in the NF1 mice were inhibited by either chemogenetic silencing of SC calcium or by systemic delivery of GDNF targeting antibodies. Together, these findings suggest that Nf1 loss in SCs causes mechanical pain by influencing adjacent neurons and, data may identify cell-specific treatment strategies to ameliorate pain in individuals with NF1.

**Graphical Abstract:** GDNF released from Schwann cells acts on sensory neurons leading to mechanical hypersensitivity and pain-like behaviors in preclinical models of NF1.

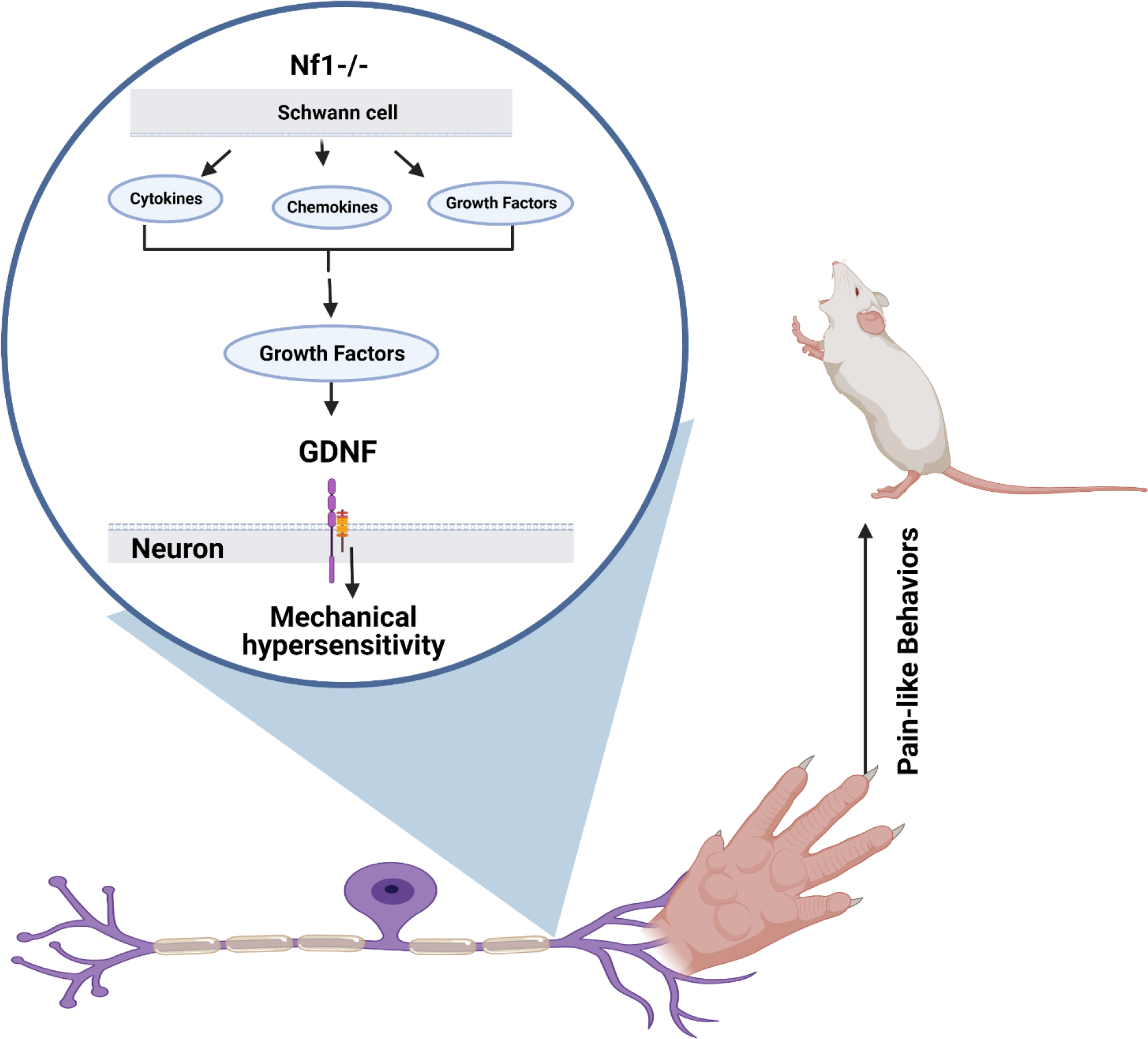

## Introduction

Primary afferent neurons transduce sensory information from the peripheral nervous system (PNS) to the central nervous system (CNS) [1–4]. Several pieces of recent evidence suggest a pivotal role for non-neuronal cells, including glia, in modulating the complex communication among cell types to regulate somatosensation [5–7]. Schwann cells (SCs) in particular have been shown to play a crucial role in nociceptive processing in the periphery. SCs are themselves mechanically sensitive and can contribute to somatosensation [5].

SC diseases often present with pain as a chief complicating factor [8, 9]. Neurofibromatosis 1 (NF1) [10] is a genetic disorder present in approximately 1/3000 live births [11, 12]. NF1 is a multisystem disorder with widespread complications which can include multiple flat, light-brown patches of skin pigmentations (café-au-lait spots), skinfold freckling, nerve tumors (cutaneous neurofibromas) under the skin, and small nodules in the iris (Lisch nodules), as well as motor and cognitive dysfunction, bone abnormalities, and predisposition to other tumor types. At least half of individuals with NF1 develop plexiform neurofibromas in the peripheral or cranial nerves which can transform to malignant peripheral nerve sheath tumors (MPNSTs) [11, 13–17]. Plexiform neurofibromas are a debilitating complication of NF1, as they can cause disfigurement and/or functional impairment. These nerve tumors present a major challenge for therapy [18] as the only curative strategy available is surgical resection, which is often not possible due to the tumor integrated nerves [19]. As a result, tumor-associated pain can be a major debilitating symptom in NF1 patients [20]. However, patient-reported pain often precedes or can be independent of tumor formation [21].

NF1 is characterized by loss of the *Nf1* gene which produces neurofibromin, a negative regulator of Ras-GTP signaling that modulates cell growth [22–24]. There are mutations in both *Nf1* alleles in neurofibromas and neurofibroma SCs [11, 25–28]. Complete *Nf1* loss of function in SCs correlates with neurofibroma formation. Therefore, SCs and/or their precursors are the known pathogenic cells in neurofibroma development [11, 25, 29]. In contrast, most non-glial cells, including sensory neurons, are wild type (in somatic mosaic patients or sporadic neurofibroma) or heterozygous for *Nf1* mutations (in most individuals with NF1) [15, 16, 30–33].

Mice that are Nf1 haploinsufficient (Nf1+/-), and a few other rodent models of NF1, have been used to model NF1-related hypersensitivity and pain; however, none of these models recapitulate all of the features of NF1 [34–37]. Genetically engineered mice that carry a homozygous deletion of *Nf1* in SCs and SC precursors causes spontaneous tumor formation over time and have become an essential tool to study NF1 tumorigenesis [38, 39]. The potential contribution of SCs lacking *Nf1* to the development of pain, however, has not been studied.

Tumors present with elevated levels of signaling molecules that include chemokines, cytokines, and various growth factors [13]. These elevated factors are known to play a prominent role in the onset of pain in many neuropathic pain-like conditions [40–42] [6, 43]. SCs may also modulate pain perception by releasing pro-algesic neurotrophic factors and cytokines/chemokines [40, 44]. While *Nf1* mutant SC express higher levels of factors than wild type cells, the contribution of SC factor release in pain development in NF1 remains unclear. Here, we found that SCs are primary contributors to hypersensitivity in a mouse model of NF1. Pain-like behaviors are observed prior to tumor formation and are regulated by enhanced GDNF expressed by SCs.

## Results

### Chemo-genetic activation of SCs induces peripheral hypersensitivity

Recent studies suggested that optogenetic stimulation of SCs can modulate nociception from the skin [40, 44]. We tested if a more physiologically relevant stimulus, enhancement of SC calcium, might induce similar alterations in afferent function and behavior. We utilized a transgenic mouse that expressed a Gq-coupled DREADD in SCs (DhhCre;hM3Dq) to allow for the artificial manipulation of SC calcium signaling. We confirmed that chemogenetic manipulation of primary SCs *in vitro* effectively increased SC calcium levels upon treatment with the designer drug, clozapine-N-oxide (CNO) [45, 46] . We also confirmed that, as expected for the Dhh-Cre driver [47], in our transgenic mice the DREADD was expressed in satellite glial cells, Schwann cells that surround putative neurons in the DRGs and in nerve Schwann cells, but not in DRG neurons (**Fig 1A**). We then performed a dose response analysis on these mice treated with CNO once daily for up to 7d to determine if SC calcium modulation might alter mechanical withdrawal thresholds *in vivo* **(Suppl. Fig 1A**). We found that delivery of CNO for seven days to 4-month-old DhhCre;hM3Dq mice *in-vivo* was sufficient to decrease mechanical withdrawal thresholds as assessed using Randall-Selitto mechanical hypersensitivity testing (**Fig 1B).** This treatment regimen also caused the animals to avoid a noxious mechanical stimulus in a mechanical conflict avoidance (MCA) assay, a more operant measure of mechanical pain (**Fig 1C**).

**Figure 1:**
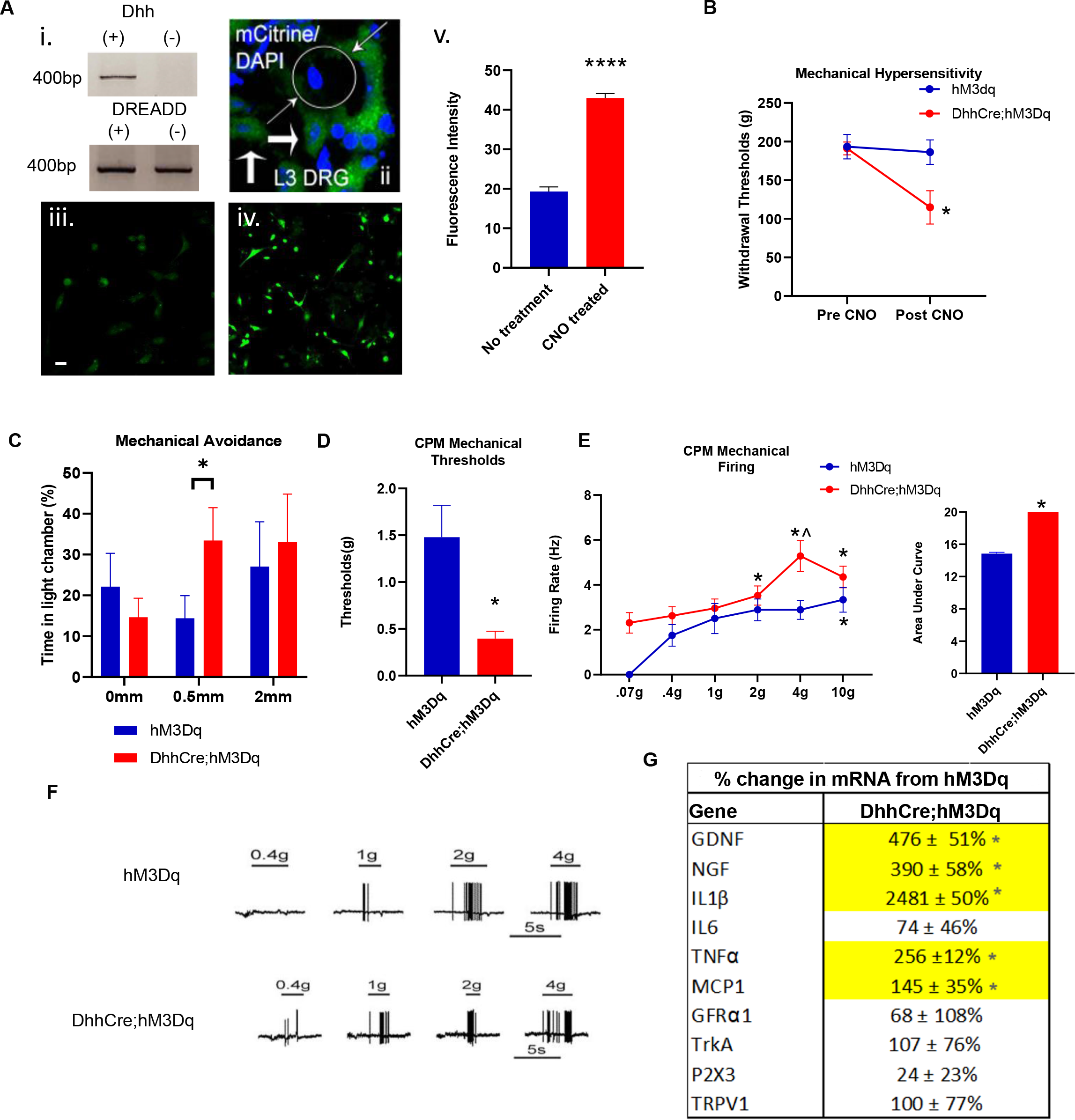
Chemogenetic activation of SCs induces peripheral hypersensitivity. **A:** DhhCre;hM3Dq (i.) mice expressing DREADD reporter mCitrine in SCs (larger arrows) surrounding putative neurons (circle with small arrows: DAPI+;mCitrine-) in DRGs (ii.). SC cultures from DhhCre;hM3Dq mice treated with CNO (40μM) display enhanced calcium fluorescence (Fluo-4) compared to untreated cultures (iii-v.) (****p <0.0001 vs no treatment, 1- way ANOVA; Mean ± SEM). **B:** 7d treatment of DhhCre;hM3Dq mice (n=5) with CNO (2mg/kg, i.p. 1x/day) induces mechanical hypersensitivity compared to CNO treated controls (n=8) by R-S (*p<0.05 vs hM3Dq post CNO, 2-way ANOVA with Tukey’s post hoc; Mean ± SEM). **C:** Similar results are also seen using the MCA assay (*p<0.05 vs hM3Dq post CNO, 1-way ANOVA with Tukey’s post hoc; Mean ± SEM). **D:** *Ex vivo* recording of saphenous afferents indicates reduced mechanical thresholds in *C-polymodal* (CPM) fibers in CNO treated DhhCre;hM3Dq mice (n=12 CPMs) compared to controls (n=12 CPMs) (*p<0.05 vs hM3Dq, 1-way ANOVA; Mean ± SEM). **E:** Enhanced firing over increasing forces is increased in the DhhCre;hM3Dq CPMs vs control CPMs. (*p<0.05 vs 0.07g, ^p<0.05 vs control 4g; 2-way RM ANOVA, HSD post hoc, Insert (AUC): *p<0.05, 1-way ANOVA; Mean ± SEM). **F:** Example firing patterns of CPM neurons from DhhCre;hM3Dq and hM3Dq mice after 7days of CNO injections. **G:** Realtime PCR analysis of L2/L3 DRGs for cytokines, growth factors and receptors/ channels indicate significant upregulation of select factors in CNO treated DhhCre;hM3Dq mice compared to controls treated with CNO (n=4/5 per group). (* and yellow highlights = p<0.05 vs controls, 1-way ANOVA with Tukey’s post hoc, Values = % change ± variance).

Changes in neuronal firing can accompany pain onset. We therefore tested if enhanced calcium in SCs might affect adjacent sensory neurons in the DRG and peripheral nerve. We used an *ex vivo* preparation which contains hairy skin, saphenous nerve, DRG, and spinal cord [48, 49]. We found that polymodal C-fibers (CPM) in the CNO treated DhhCre;hM3Dq mice were sensitized to mechanical but not heat stimulation of their receptive fields compared to CNO treated controls. Mechanical thresholds were reduced (**Fig. 1D**), while firing rates to mechanical stimuli over the range of forces tested was increased in DhhCre;hM3Dq CPMs compared to control CPMs (**Fig 1E-F**). Effects of SC-mediated sensitization appeared to be specific to the CPM neuron subpopulation (**Suppl. Fig. 1B-F**).

Previous work has shown that SCs are sources of a variety of growth factors and cytokines that are considered pro-algesic, and *Nf1* mutant Schwann cells produce increased levels of such factors [40, 50–52]. We performed a small screen of factors known to be produced by SCs in the DRGs of the DhhCre;hM3Dq mice treated with CNO. We found that enhancing calcium in SCs caused a significant upregulation of mRNAs encoding several growth factors and cytokines that could affect peripheral sensitivity including glial cell line-derived neurotrophic factor (GDNF), nerve growth factor (NGF), interleukin 1β (IL1β), tumor necrosis factor α (TNFα) and the GDNF co-receptor, GFRα1 (**Fig 1G**). Together these data suggest that alterations in SCs might be partially responsible for altering specific sensory neurons that modulate mechanical responsiveness.

### Deletion of *Nf1* in SCs but *NOT* sensory neurons causes mechanical hypersensitivity

Our data suggested that SCs play an important role in sensory processing in the periphery. We therefore determined if in a disease model with altered SC biology, whether altered SCs contribute to pain. We focused on NF1, as pain is a debilitating symptom in NF1 patients [18, 53], and is not always associated with nerve tumors [20]. To determine if *Nf1* deletion in neurons and/or SCs causes hypersensitivity, we evaluated behavioral responsiveness in a sensory neuron *Nf1* mutant mouse (PirtCre;Nf1^+/f^) and in a SC specific *Nf1* mutant (DhhCre;Nf1^f/f^). Use of these Cre lines coordinated the timing of deleting *Nf1* from neurons and SC to ∼E11-12 [13, 47]. To recapitulate cell mutational status in most individuals with NF1 [47, 54] we also assayed mice with heterozygous deletion in sensory neurons (PirtCre;Nf1^+/f^) and homozygous deletion in SCs (DhhCre;Nf1^f/f^). Finally, we used mice in which neurons, Schwann cells, and all other cell types are haplo-insufficient for *Nf1* (i.e., Nf1+/-). Using standard evoked cutaneous mechanical hypersensitivity assays (Randall Sillito) on the hairy skin of the hind paw [55], similar to previous reports in uninjured mice [56], haplo-insufficient knock-out mice showed no significant difference in responsiveness over time when compared to littermate controls (**Fig 2A).** Mice with a copy of *Nf1* deleted in sensory neurons (PirtCre;Nf1^+/f^) also did not show mechanical hypersensitivity by Randall-Sillito at any time point (**Fig 2B**). DhhCre;Nf1^f/f^ mice [47] again did not display mechanical hypersensitivity at any time points tested compared to controls (**Fig 2C).**

**Figure 2:**
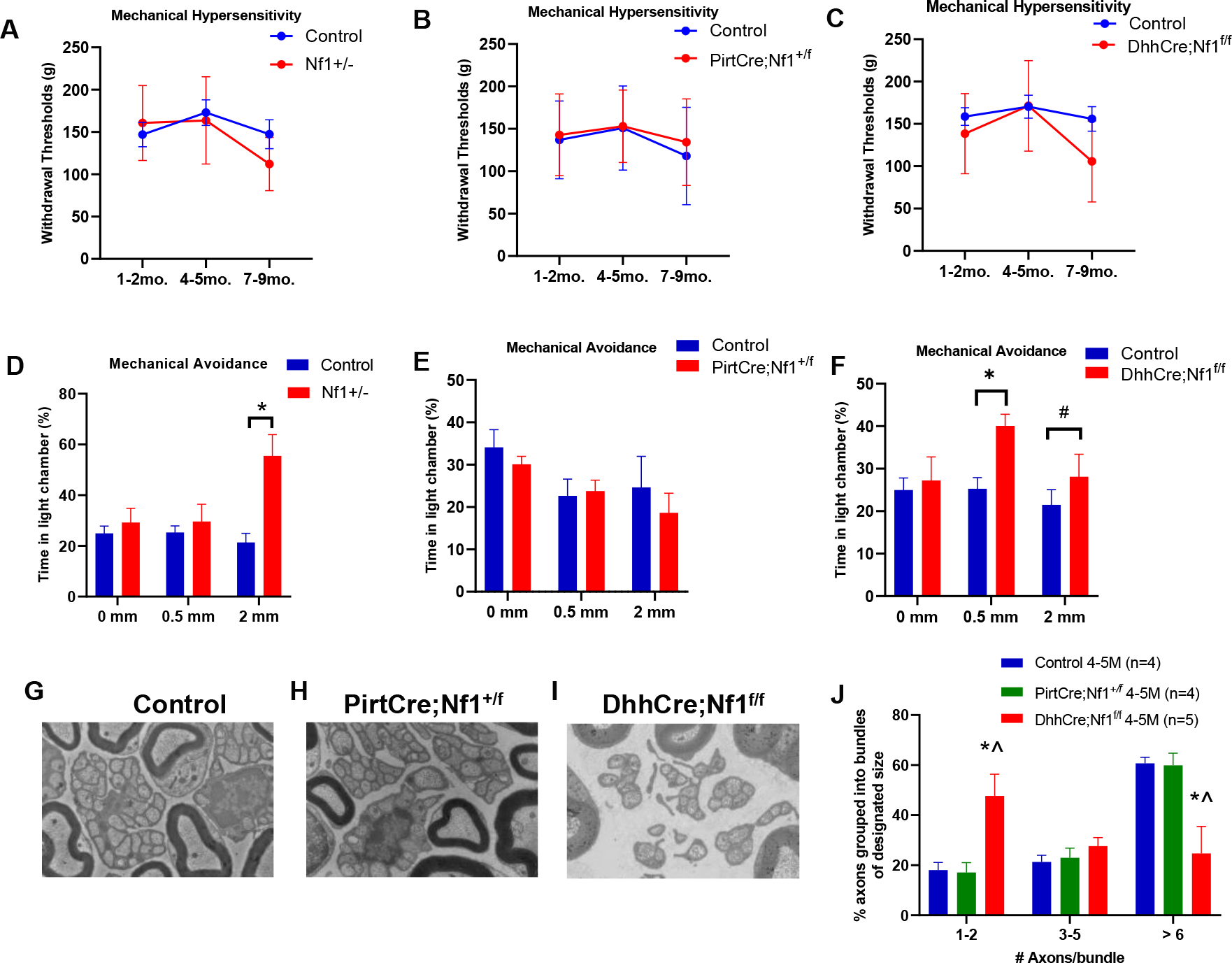
**Schwann cell specific knockout of *Nf1* leads to mechanical hypersensitivity**. **A**: *Nf1* haploinsufficient mice (Nf1+/-) do not show any mechanical hypersensitivity using the Randall-Selitto assay (n=12 control; n=13 mutant). **B:** The deletion of *Nf1* in sensory neurons (PirtCre;Nf1^+/f^ ) also didn’t show any mechanical hypersensitivity using R-S (PirtCre;Nf1^+/f^; n=17 control n=17 mutant). **C**: Mice with SC specific *Nf1* deletion (DhhCre;Nf1^f/f^;) show a modest reduction in mechanical withdrawal threshold (n=20 control; n=12 mutant; p>0.05 vs time matched littermate controls; 2-way RM ANOVA, Tukey’s post hoc; Mean ± SEM). **D:** Using MCA assay, Nf1+/- mice prefer to spend more time during the assay exposed to an aversive light stimulus compared to a noxious mechanical stimulus. (*p<0.05 vs controls, 1-way ANOVA with Tukey’s post hoc; Mean ± SEM). **E**: PirtCre;Nf1^+/f^ mice did not show any significant difference in time spent in either light or dark chambers. **F:** DhhCre;Nf1^f/f^ mice display increased mechanical avoidance even with smaller spikes present vs. littermate controls. (*p<0.04, #p<0.054 vs. controls 1-way ANOVA with Tukey’s post hoc; Mean ± SEM). **G**: Electron micrograph of the saphenous nerve shows intact Remak bundles surrounding the axons in Nf1+/- mice at 4-5 months (Control). **H**: Representative electron micrograph of saphenous nerves in PirtCre;Nf1^+/f^ shows intact Remak bundles at 4-5 months. **I**: However, significant Remak bundle disruption is observed in the saphenous nerve of 4-5-month-old DhhCre;Nf1^f/f^ mice. **J**: Quantification of percentage of axons grouped within a Remak bundle in the groups (*p<0.05 vs control; ^p<0.05 vs PirtCre;Nf1^+/f^, 2-way ANOVA with Tukey’s post hoc; Mean ± SEM).

We then tested animals in a more operant task, the mechanical conflict avoidance (MCA) assay [57, 58]. This allows the animal to freely choose between aversive stimuli. In the MCA test, 4-month-old Nf1+/- animals displayed enhanced mechanical avoidance indicating that this assay is sensitive for assessing pain-like behaviors in models of NF1 (**Fig 2D**). However, when using the MCA assay in 4-month-old PirtCre;Nf1^+/f^ mice, no differences in mechanical hypersensitivity were observed compared to controls (**Fig. 2E**). Combining the heterozygous deletion of *Nf1* in both sensory neurons and SCs did not alter mechanical avoidance in the MCA assay **(Suppl. Fig 2A),** suggesting that other cell types might contribute to effects in Nf1+/- mice. Importantly, DhhCre;Nf1^f/f^ mice at 4-months did show increased mechanical avoidance (**Fig. 2F).** Of note, when assessing intercrossed PirtCre;Nf1^+/f^ and Nf1^f/f^ mice, only 1/37 animals from these litters had both copies of *Nf1* deleted from sensory neurons (PirtCre;Nf1^f/f^), suggesting issues with survivability with complete deletion of *Nf1* from primary afferents at embryonic stages.

Since previous work has shown that deletion of *Nf1* in SCs causes disruptions in nerve structure, we also assessed Remak bundle integrity in our groups at 4-5 months of age, a time when tumors are not yet present in the lumbar DRGs or saphenous nerves [47]. The saphenous nerve from Nf1+/- and PirtCre;Nf1^+/f^ mice displayed no significant alterations in Remak bundle structure (**Fig. 2G-K**) at 4-5 months. In the DhhCre;Nf1^f/f^ mice, the disruption of the Remak bundles increased significantly during this time frame. Together this suggests that Schwann cells play an important role in the onset of hypersensitivity in NF1 prior to tumor formation, but during time periods of Remak bundle disruption.

### Mechanical but not thermal hyper-responsiveness is observed in primary afferents of mice with SC deletion of *Nf1*

Our behavioral data suggested that SCs are key players in NF1 related hypersensitivity. Therefore, we determined if deleting *Nf1* from SCs alters primary afferent responsiveness, using *ex vivo* recording (**Fig 3A-B).** We found that in the DhhCre;Nf1^f/f^ mice, the myelinated, high threshold mechanoreceptors (HTMRs) displayed a significant reduction in mechanical thresholds (**Fig. 3C**) and an increase in firing to mechanical stimulation of their receptive fields compared to wildtype (WT) C57Bl/6 and Nf1^f/f^ controls (Cre negative) but showed no change in heat responsiveness (**Fig 3D-E**). Polymodal C-fibers (CPM) in the DhhCre;Nf1^f/f^ mice also displayed reduced mechanical thresholds and enhanced firing rates in response to mechanical stimuli, but no change in heat sensitivity compared to controls (**Fig 3F-H**). No changes in response properties were observed in other neuronal subtypes between groups (**Suppl. Fig. 3**). These results suggest that SCs play a role in the sensitization of adjacent sensory neurons to mechanical stimuli, which could underlie pain-like behaviors.

**Fig 3:**
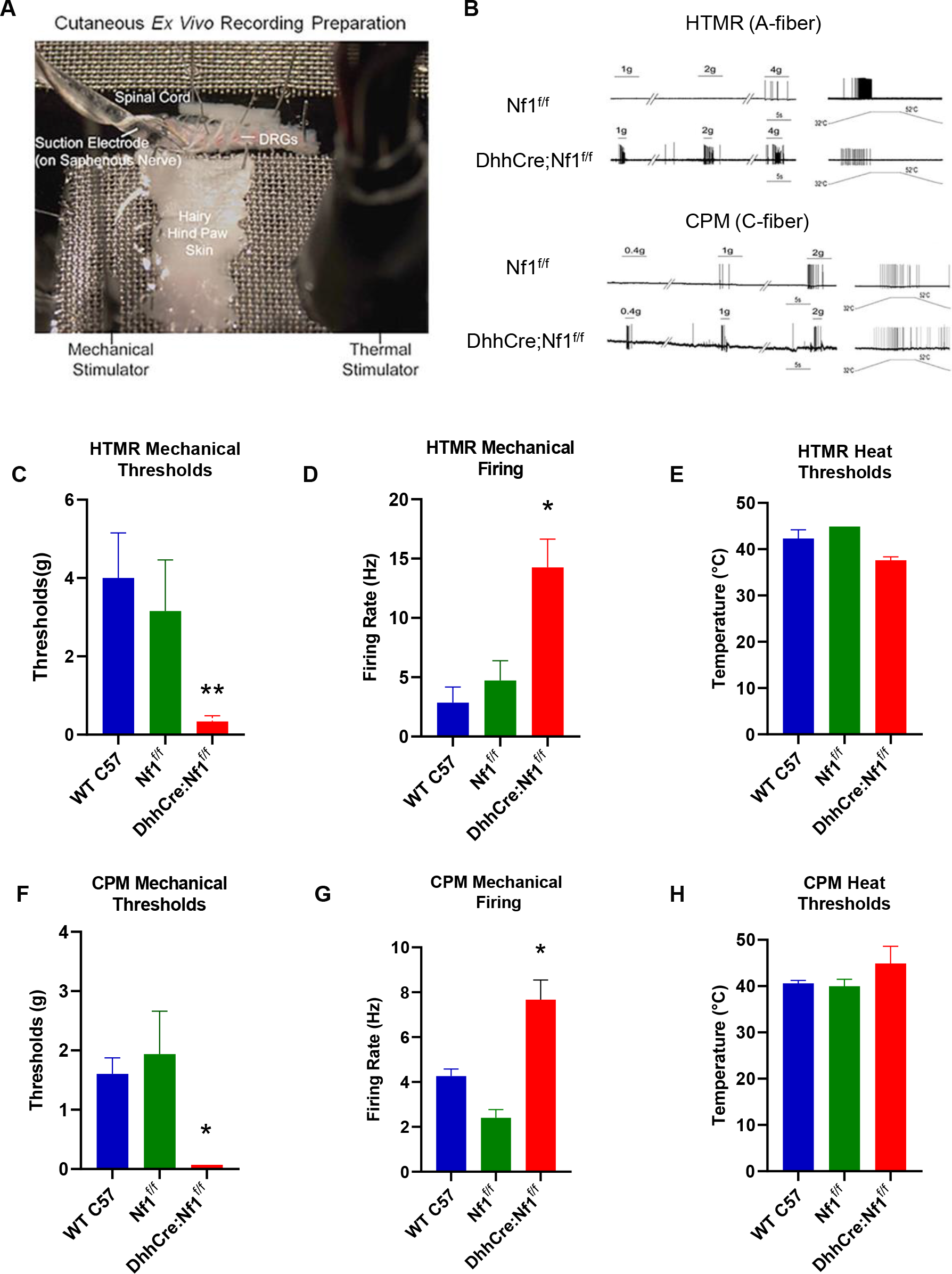
Sensitization of high threshold mechanoreceptors (HTMRs) and polymodal C-fibers (CPM) in DhhCre;Nf1^f/f^ mice as assessed with *ex-vivo* recording. **A:** Representative image of the *ex vivo* electrophysiological recording preparation. **B:** Firing pattern of A-fiber HTMRs and polymodal C-fibers (CPM) in DhhCre;Nf1^f/f^ mice at 4-5 mo of age. **C**: HTMRs showed a significant reduction in mechanical thresholds compared to control HTMRs (WT C57, n= 9; Nf1^f/f^, n=9, DhhCre;Nf1^f/f^, n=7; **p<0.01 vs. WT C57 and Nf1^f/f^, 1-way ANOVA with Tukey’s post hoc; Mean ± SEM; Total no. of cells, WT C57, n=45; Nf1^f/f^, n=57, DhhCre;Nf1^f/f^, n=37). **D**: Firing rates of HTMRs showed the increased firing to mechanical stimuli in DhhCre;Nf1^f/f^ mice when compared with controls (*p<0.05 vs. WT C57 and Nf1^f/f^, 1-way ANOVA with Tukey’s post hoc; Mean ± SEM). **E**: HTMRs showed no change in heat thresholds. **F**. Polymodal C-fibers (CPM) in DhhCre;Nf1^f/f^ mice also showed reduced mechanical thresholds compared with controls (WT C57, n= 14; Nf1^f/f^, n=14, DhhCre;Nf1^f/f^, n=8; *p<0.05 vs. WT C57 and Nf1^f/f^, 1-way ANOVA with Tukey’s post hoc; Mean ± SEM). **G:** DhhCre;Nf1^f/f^ mice also showed the increased firing rate of CPMs (*p<0.05 vs. WT C57 and Nf1^f/f^, 1-way ANOVA with Tukey’s post hoc; Mean ± SEM). **H**: CPMs in DhhCre;Nf1^f/f^ mice showed no change in heat thresholds.

### Inhibition of enhanced SC calcium in DhhCre;Nf1^f/f^;hM4Di mice reduces mechanical hypersensitivity

Previous work has shown that Nf1-/- SC have significantly elevated calcium responses to stimulation with ATP [59]. ATP acts through GPCRs on the SC surface, which couple to changes in calcium via activation of downstream signaling though small g proteins. We tested if we could reverse mechanical hypersensitivity observed in the NF1 mouse model by use of inhibitory DREADDs in SC (DhhCre;Nf1^f/f^;hM4Di), which will suppress calcium intracellularly [60]. We confirmed that SC isolated from DhhCre;Nf1^f/f^;hM4Di mice display enhanced calcium responses to ATP stimulation. Treatment of SC cultures with the DREADD agonist C21 significantly inhibited the ATP-induced calcium response (**Fig 4A-B**). C21 was used in these experiments in order to avoid non-specific effects of high dose CNO, which are often required for activation of the inhibitory DREADD *in vivo* [59]. In the MCA assay, prior to C21 treatment, we found that DhhCre;Nf1^f/f^;hM4Di mice displayed the expected increase in mechanical avoidance. However, after treating these mice with C21 for 7d, even without any noxious mechanical stimulus added, mice spent an equal amount of time in both light and dark chambers indicating that inhibition of SC may affect light sensitivity (**Suppl. Fig. 4**). We therefore modified this assay to avoid the use of light as an aversive stimulus. Instead, we allowed mice to perform the task when all chambers are dark; we provided one side with home cage bedding. Normal mice choose to spend more time in the home-bedding chamber in this assay; however, DhhCre;Nf1^f/f^;hM4Di mice, prior to C21 treatment, spent less time crossing the noxious mechanical stimulus in order to reach the home-bedding chamber. After 7d of C21 treatment, however, these same mice showed no differences compared to controls (**Fig. 4C**). These results strongly suggest that that enhanced calcium signaling in SC is a major driver of pain-like behaviors in this mouse model of NF1.

**Figure 4:**
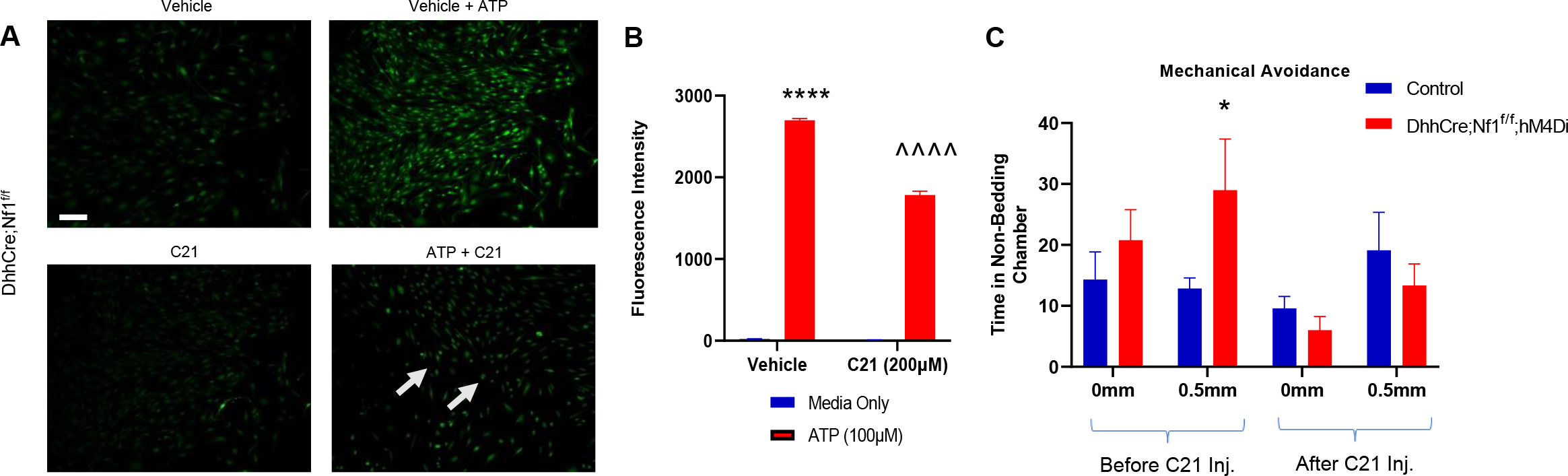
Chemogenetic inhibition of SCs suppresses mechanical hypersensitivity in DhhCre;Nf1^f/f^;hM4Di mice. A: In isolated Schwann cells from sciatic nerves of DhhCre;Nf1^f/f^;hM4Di mice, no significant changes in calcium are detected in SCs treated with vehicle (DMSO). Calcium release is increased upon addition of ATP (100µM with Vehicle). No significant changes in calcium are detected in SCs treated with compound 21 (C21) alone. Inhibition of ATP-induced calcium is observed however with C21 in SCs isolated from DhhCre;Nf1^f/f^;hM4Di mice. **B:** Quantification of fluorescence intensity from SCs depicting changes in calcium release from conditions outlined in A (****p<0.0001 ATP with vehicle vs ATP with C21 and ^^^^p< 0.0001 C21 vs ATP with C21, 2-way ANOVA with HSD post hoc; Mean ± SEM). **C**: DhhCre;Nf1^f/f^;hM4Di mice display increased mechanical avoidance even with smaller spikes present vs. littermate controls (n=16 control, n=7 mutant, *p <0.05 vs controls, 2-way ANOVA, Tukey’s post hoc; Mean ± SEM), before C21 injection but after 7 days of C21 injection (i.p.), mechanical avoidance is reduced to control levels

### SC specific deletion of *Nf1* alters gene expression in the DRGs

Increased levels of cytokines, growth factors and other molecules have been found in neurofibromas of DhhCre;Nf1^f/f^ mice [13], and in Schwann cells derived from these mice. Many of these molecules are known to play important roles in the modulation of pain [51, 52, 61–63]. To begin to determine mechanisms through which SCs cause peripheral sensitization under normal and pathological conditions, we performed additional analysis of existing single cell RNA-Seq data obtained from the DRGs of DhhCre;Nf1^f/f^ mice and controls at 2 months of age [64, 65]. Of the cytokine/chemokine/ growth factor transcripts that differed in SC clusters between control and Nf1 mutants, GDNF was the only factor upregulated in SC precursors and in non-myelinating SCs (**Fig. 5A**). We also used the CellChat algorithm to predict cell types that express GDNF receptors; DRG neuron types were identified based on Usoskin et al [66]. This signaling prediction analysis indicated that enhanced SC derived GDNF targets a variety of sensory neuron subtypes including the non-peptidergic neurons (**Fig. 5B**) that are likely those observed to be sensitized in the DhhCre;Nf1^f/f^ mice as defined by *ex vivo* recording (**see Fig. 3**). Other genes deregulated in SC and other cell types that may influence neurons are shown in (**Suppl. Fig. 5A-C).** qRT-PCR validated that levels of GDNF transcript (*p<0.05 vs Nf1^f/f^ controls; 1-way ANOVA) were selectively elevated in DhhCre;Nf1^f/f^ DRGs compared to controls (**Fig. 5C).** In contrast, realtime PCR showed that genes were differentially altered as a result of SC/SCP specific loss of *Nf1*, were not affected by sensory neuron *Nf1* loss (**Suppl. Fig. 5D**).

**Fig 5:**
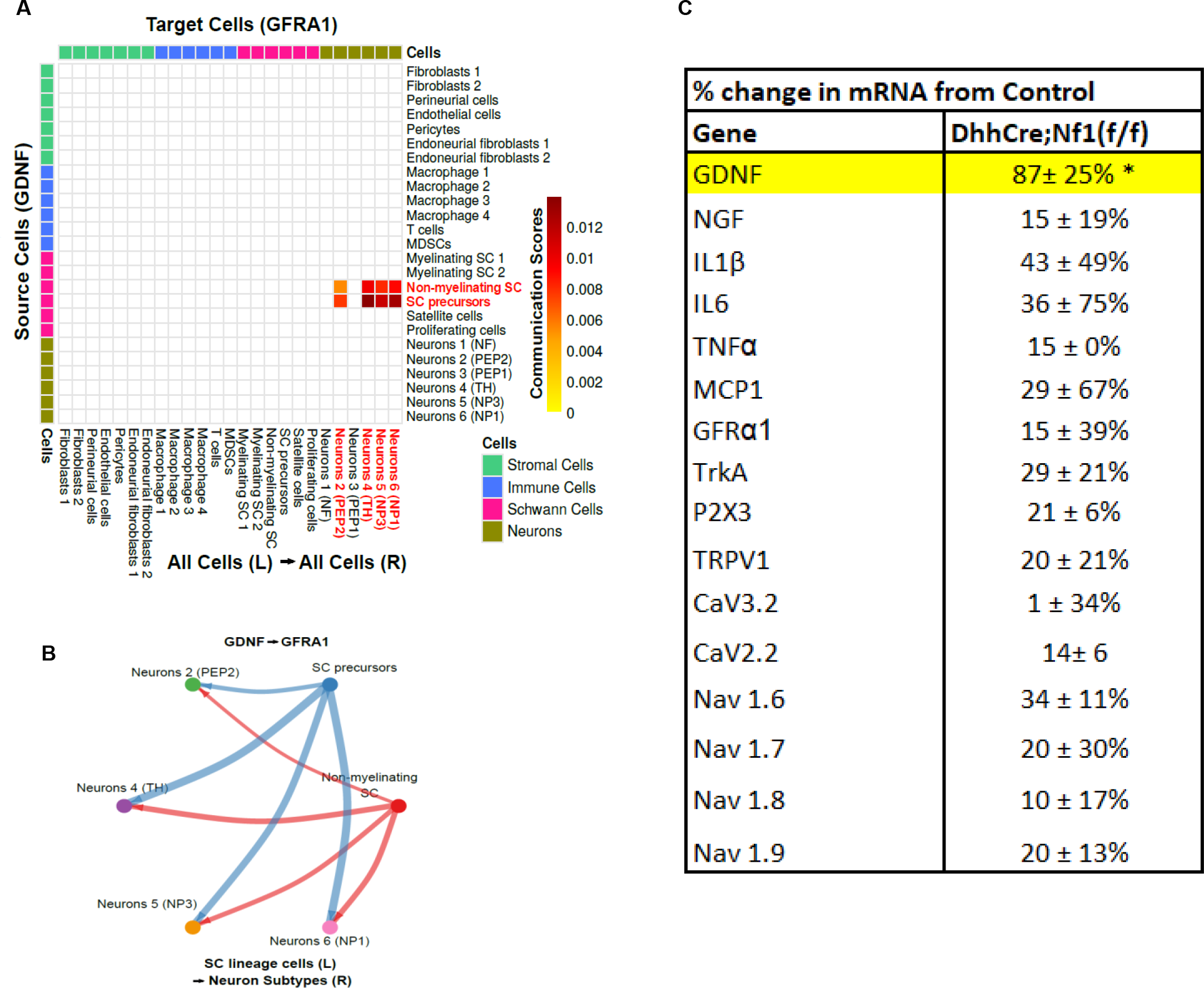
GDNF signaling from SC to neurons is enhanced in DhhCre;Nf1^f/f^ DRGs. A: Analysis of cell-cell signaling in DhhCre;Nf1^f/f^ DRGs indicates that GDNF signals from non-myelinating SC and SC precursors to neurons containing its co-receptor (GFRα1). **B:** Four neuronal subtypes (TH+, peptidergic 2 (PEP2), non-peptidergic 3 (NP3) and NP6) display unique signaling of GDNF with SC precursors (SCP) and non-myelinating SC in 7 mo tumor compared to 2 mo control/pretumor or 7 mo control. **C:** Realtime PCR from DhhCre;Nf1^f/f^ DRGs shows elevated GDNF expression, validating scRNA-Seq (* and yellow highlights = p<0.05 vs controls, 1-way ANOVA with Tukey’s post hoc, Values = % change ± variance).

We then confirmed the increased expression of GDNF in SCs of the DRG and quantified subpopulations of sensory neurons using immunohistochemical analysis. We found a significant increase in GDNF in S100β+ satellite glial cells and SC of the DRG in DhhCre;Nf1^f/f^ mice compared to controls (**Fig 6 A-B).** No changes in the neuronal markers TRPV1, IB4 or ASIC3 were found in the DRGs of DhhCre;Nf1^f/f^ mice compared to controls (**Suppl. Fig. 6**). To determine if SC produced GDNF plays a role in the hypersensitivity in the NF1 mouse model, we treated DhhCre;Nf1^f/f^ mice *in vivo* with a GDNF-targeting antibody and performed MCA analysis. Up to 48 hours after injecting DhhCre;Nf1^f/f^ mice with a function-blocking GDNF antibody, the mechanical hypersensitivity that is normally observed in these animals was no longer observed (**Fig. 6C; Suppl. Fig. 6**).

**Fig 6:**
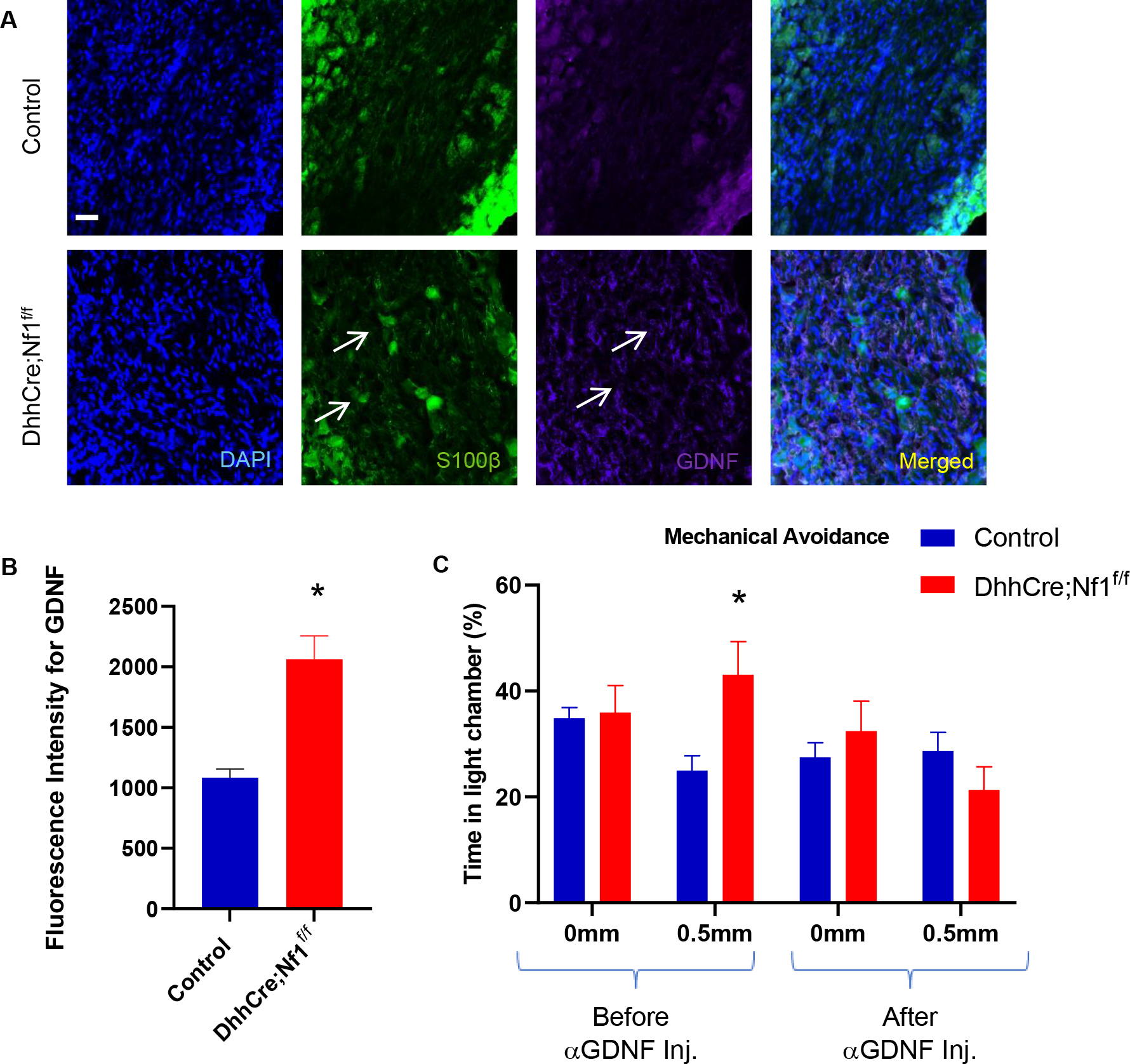
GDNF is elevated in the dorsal root ganglion of DhhCre;Nf1^f/f^ when compared to control mice and regulates behavioral hypersensitivity. A. Representative images of DRGs stained with different markers including S100b (green), GDNF (purple) and DAPI (blue). GDNF is mainly expressed in glial cells (arrows). Scale bar, 50µm. **B.** Quantifying the fluorescence intensity from each image shows elevated GDNF in DRGs of DhhCre;Nf1^f/f^ mice compared to controls (*p<0.05 vs. control, 1-way ANOVA with Tukey’s post hoc; Mean ± SEM). **C.** DhhCre;Nf1^f/f^ mice normally display mechanical hypersensitivity in the MCA assay at 4 months; however, 24hrs after being injected (i.v.) with GDNF targeting antibody, mechanical avoidance is reduced to control levels. (*p < 0.05 vs controls, 2-way ANOVA, with Tukey’s post hoc; Mean ± SEM).

## Discussion

Our data confirm an important role for SC in nociceptive processing. The targeted chemogenetic activation of Schwann cell calcium caused increased hypersensitivity at the afferent and behavioral levels that correlated to altered DRG gene expression (**Fig. 1**). Similar results were observed when knocking out the *Nf1* gene in SC/SCPs (but not sensory neurons), prior to tumor formation, in a genetically engineered mouse model of NF1 [47] (**Fig. 2-3**). Blocking enhanced SC calcium in the DhhCre;Nf1^f/f^ mice using inhibitory DREADDs blunted the observed mechanical hypersensitivity (**Fig. 4**). Of the factors upregulated in DRGs by DREADD dependent activation of SC/SCPs, and by specific deletion of *Nf1* in these cells, we identified distinct induction of GDNF expression, which corresponded with enhancement of GDNF signaling to neurons from SCs as assessed by scRNA-Seq analysis (**Fig. 5**). Finally, targeting GDNF with systemic antibody treatment reduced mechanical hypersensitivity in the NF1 mouse model (**Fig. 6**).

Glial cells play a pivotal role in the functioning of the nervous system. Multiple roles from regulating neuronal survival and differentiation during embryogenesis [67], to modulating the formation of myelin sheaths, maintaining the appropriate concentrations of ions in the nerve mileu, and regulating nociception are known [5, 6, 67]. In the periphery, Schwann cells are known to provide the first response to nerve injury to initiate repair and facilitate axon regeneration [68]. Recently, Schwann cells have been shown to also play a role in the development and maintenance of pain by proliferating and interacting with nociceptive neurons to release factors such as chemokines/cytokines/growth factors [5, 7, 69, 70]. As recent studies have been focused on neuron-glia cross talk, strategies targeting this interaction have gained traction as potential therapies for pain.

Here we found that DREADD-dependent activation of SCs increased SC calcium signaling, which was alone sufficient to induce mechanical hypersensitivity in adjacent sensory neurons. This afferent sensitization likely underlies the behavioral hypersensitivity found in these transgenic mice (**Fig. 1**) [4, 44, 71–73]. We also found that chemogenetic activation of SCs could upregulate a specific set of growth factors and cytokines that may influence sensory function (**Fig. 1**). There are, of course, a number of additional ways that Gq signaling in SCs could alter somatosensory processing. These include regulation of ion channels that can modulate the electrochemical gradient in the nerve and/or release of other factors that may modify structural integrity of the nerve [74–77]. These possibilities will need to be explored in future studies.

Previous work on NF1-related pain has focused on use of haploinsufficient mice (Nf1+/-), editing of *Nf1* in adult animals using gRNAs, or studied the release of the different neuropeptides from sensory neurons of the Nf1+/- animals under injury conditions [21, 78]. The *in vivo* studies have provided some information on how pain may develop in NF1, but they do not provide understanding of how specific cell types contribute to the onset of NF1 associated pain, because in both haploinsufficient mice and after gene editing, multiple nerve cell types are affected. Studies using dissociated neurons *in vitro* have suggested that these cells can display enhanced excitability upon *Nf1* mutation [79] but *in vivo*, an optimized environment may be necessary to observe sensitization (**Figs. 2,3**).

Prior studies also did not address the timing of pain onset [21, 26, 34, 35]. We therefore utilized several transgenic lines define cell types involved in hypersensitivity upon *Nf1* mutation. When performing standard evoked hypersensitivity assays on *Nf1* haplo-insufficient mice, minimal effects are seen over time, consistent with previous work [34]. However, by using behavioral tests that provide a choice for the animal, such as the MCA, Nf1+/- mice show mechanical hypersensitivity (**Fig. 2**). Similar to the Nf1+/- mouse, sensory neuron *Nf1* mutants and SC/SCP *Nf1* mutants display minimal effects using R-S testing. However, MCA analysis reveals a role for SC/SCP *Nf1* in pain-like behaviors that is not observed in the sensory neuron mutants (**Fig. 2**). This indicates that assays that allow the animal to choose between stimuli may be more sensitive for pain analyses in models of NF1. DhhCre;Nf1^f/f^ mice show afferent mechanical sensitization (**Fig 3).** In contrast, the PirtCre;DhhCre;Nf1^+/f^ mice **(SF 2)** do not show any alterations in mechanical responsiveness, suggesting a prominent role for SCs in pain development in NF1.

In the DhhCre;Nf1^f/f^ preclinical model of NF1, tumors form in the cervical region around 4 months of age. Small tumors can form in the lumbar DRG, which innervate the hindlimb; however, tumors are not visible until 6-9 months. Tumors are preceded by disruptions in nerve structural integrity that are also present in human nerve and tumors [39, 80]. Although we cannot rule out a role for Remak bundle disruption (**Fig. 2**) in pain-related behaviors in the DhhCre;Nf1^f/f^ mice, results from the GqDREAAD experiments (**Fig. 1**) indicate that SCs are capable of inducing hypersensitivity alone. Future experiments will be needed to address how or if Remak bundle disruption contributes to pain in NF1.

In our choice assay, DhhCre;Nf1^f/f^ mice with chemogenetic inhibition of SC calcium (e.g. suppression of the enhanced calcium found in Nf1-/- SCs [80]) reversed observed mechanical hypersensitivity (**Fig. 4**). However, it was important in this assay to eliminate light as an aversive stimulus (**Suppl. Fig. 4**). Previous reports in other models of NF1 suggest that these mice may display enhanced light sensitivity [81]. Chemogenetic inhibition of calcium in SC/SCPs mutant for *Nf1* appeared to have resulted in a loss of light aversion. It will be important in the future to assess light sensitivity in these various mouse models of NF1. Another point to note is that the GiDREADD can also affect cAMP and potassium efflux in addition to calcium [45]. It will be necessary in future experiments to determine the potential roles, if any, of these other factors in NF1-related hypersensitivity. In spite of these limitations, our data indicate that prior to tumor formation, mutations in SC/SCP *Nf1* are key players in pain-like behaviors. This corelates with clinical reports that individuals with NF1 often report pain in parts of the body that are not obviously affected by tumors [20].

SC/SCP deletion of *Nf1* induced robust mechanical sensitization in HTMRs and CPM neurons that could underly behavioral hypersensitivity (**Fig. 3**). Intriguingly, heat hypersensitivity was not observed, consistent with some models of NF1 as well as patient reports of a lack of heat-related pain [34]. Our results are not consistent with recent reports in which intrathecal injections of guide RNAs targeted *Nf1* in adult rats [21], possibly because that deletion was targeted to the adult nervous system whereas our model deletes *Nf1* specifically in SC/SCPs beginning around E12.5 [39]. Our results are also inconsistent with reports that show heat hypersensitivity in the Nf1+/- mouse after injury [14]. The enhanced heat hypersensitivity observed after injury in the Nf1+/- mice could indicate that this hypersensitivity can be observed if the environment is optimized for Nf1+/- Schwann cells, for example when immune cells are recruited to the nerve after injury. This explanation is consistent with the increase in tumor formation in NF1 mice after nerve injury [82, 83].

SCs can modulate nociception by releasing different factors including chemokines, growth factors and cytokines [7, 10, 42, 50, 51, 63, 84–86]. Neurotrophic factors enable neuronal outgrowth, and alterations in levels of these factors can influence peripheral sensitization [42, 48, 87, 88]. An intriguing finding in our study was that there was no increase in cytokines/growth factors in the DRGs from mice with sensory neuron *Nf1* KO (PirtCre;Nf1^+/f^) **(SF 5D)**. Rather, GDNF was elevated uniquely in the SC/SCP *Nf1* knockout (**Fig 5C**). Further, pathway analyses using scRNA-Seq data from neurofibromas indicated that of all signaling pathways that are predicted to be increased in non-myelinating SCs and Schwann cell precursor-like cells for communication with neurons, GDNF signaling was the only one specifically elevated (**Fig. 5**). Both GDNF and the related GDNF family factor, artemin, have been linked to afferent sensitization and pain in animal models and in clinical studies, so that targeting this signaling molecule has gained interest as a therapeutic strategy for pain [87, 89]. After confirming GDNF expression in glial cells of the DhhCre;Nf1^f/f^ PNS (**Fig 6**), we found that treatment of DhhCre;Nf1^f/f^ mice with GDNF targeting antibodies suppressed noxious mechanical avoidance in the MCA assay for at least 48hrs (**Fig. 6**). This result strongly supports a major role of Schwann cells in modulating afferent sensitization in NF1. This concept is further supported by our finding that non-peptidergic CPM neurons were predicted to be affected by GDNF in the DhhCre;Nf1^f/f^ mice. These cells are known to be IB4+ and GFRα1+ [90], and directly respond to GDNF. Together, our findings contribute to the increasing pool of evidence that implicates interactions between non-neuronal cells and sensory neurons in effects on nociception and extends it by application to NF1.

Pain can significantly impede daily activities in NF1 patients, yet treatment for pain in NF1 remains a significant a challenge for clinicians [21, 34–36]. SC/SCPs have been well established to play an important role in tumor formation [13, 39, 91] and our data herein suggests they also play a significant role in pain-like behavior independent of tumors. This study therefore suggests a novel approach to treat pain in NF1.

## Author Contributions

NGRR contributed to experimental design, conducted the majority of the experiments, performed data analysis and prepared and edited the manuscript. LAM performed ex vivo recording experiments on DhhCre;hM3Dq mice. LMO performed PCR and behavioral analyses on select cohorts. IM performed PCR, ex vivo recording and some behavioral tests. AA contributed to conducting cell culture experiments, data analysis and interpretation for ex vivo recordings, and treatment with DREADD agonist (compound 21). KLS contributed to IHC and PCR experiments. ARR performed additional IHC and PCR tests along with data analysis. LB performed supplementary MCA assays and PCR. MCH performed all genotyping and animal husbandry. JPC contributed to cell culture experiments and confocal imaging. TAR performed electron microscopy (EM) and data analysis. LFQ contributed to experimental design and performed behavioral and electrophysiological tests along with manuscript editing. KC performed scRNA-Seq, data analyses and figure generation. NR contributed to experimental design, provided transgenic animals, oversaw EM studies and sc-RNA- seq analyses and edited the manuscript. MPJ contributed to experimental design, performed ex vivo recordings, contributed to confocal imaging, analyzed and interpretated the data, and prepared the manuscript.

## Methods

### Animals

Male and female mice between 1 and 7 months of age were used in all studies. Mice expressing a Gq-coupled designer receptor exclusively activated by designer drugs (DREADD) specifically in SCs were used in initial experiments. To generate this mouse, we used the desert hedgehog (Dhh)-Cre mouse which expresses Cre recombinase in SCs and SC precursors. This line was crossed to a Cre-dependent Gq-coupled DREADD mouse (Rosa26-LSL-hM3Dq) to obtain a line that allows for DREADD dependent modulation of SC activity. In other studies, to knockout *Nf1* in SCs and SC precursors, we crossed the Dhh-Cre mouse to a neurofibromin 1 (*Nf1*) floxed (Nf1^f/f^) line to create the DhhCre;Nf1^f/f^ mouse model of NF1 [47]. Similarly, to target deletion of *Nf1* to sensory neurons, we utilized the PirtCre mouse (generously donated by Dr Xinzhong Dong) which targets Cre recombinase expression in sensory neurons (Kim et al 2008) and crossed it with the Nf1^f/f^ mice. Nf1+/- haplo-insufficient mice [34] and mice with SC and sensory neuron heterozygous mutations in *Nf1* (DhhCre;PirtCre;Nf1^+/f^) were used for comparisons. Additional experiments were performed as indicated on DhhCre;Nf1^f/f^ mice that contained a Cre-dependent Gi-coupled DREADD (hM4Di) in SCs. Mice were housed in a barrier facility that were maintained on a 14:10 hr light–dark cycle with a temperature-controlled environment and given food and water ad libitum. All procedures were approved by the Cincinnati Children’s Hospital Institutional Animal Care and Use Committee and adhered to NIH Standards of Animal Care and Use under the Association for Assessment and Accreditation of Laboratory Animal Care International (AAALAC)-approved practices.

### Treatments

Mice were treated with DREADD agonist clozapine-N-oxide (CNO) at 2mg/kg/d for 1-7d or compound 21 (C21) at 20µg/µL/day for 1-7d *in vivo* along with their littermate controls. In other experiments, mice were injected with GDNF-targeting antibody intravenously at 5µg/g (#ANT-014, Alomone) *in vivo* along with their littermate controls. For dissociated Schwann cell experiments *in vitro*, cells were treated with CNO at 10-40μM alone or C21 at 20-200µM with or without 100 µM of adenosine triphosphate (ATP).

### Pain-Related Behaviors

To assess evoked hypersensitivity, nociceptive withdrawal thresholds were determined using a Randall-Selitto apparatus. Before the test, the animal was acclimatized in the behavior room for 25-30 minutes. The animal was scuffed, carefully immobilized, and the right paw was placed on the platform with an application of an increasing mechanical force, in which the tip of the device was applied onto the medial portion of the hairy skin surface of the hind paw until a withdrawal response was observed. The maximum force applied was limited to 250 g to avoid skin damage. The test was repeated 3X with a 5-minute interval between stimuli [55]. The average of the three trials was determined per mouse and data was averaged per condition for comparisons.

To assess the animals’ choice to avoid either an aversive light stimulus or a noxious mechanical stimulus, the mechanical conflict avoidance (MCA) assay was used [92]. Mice were placed in a chamber for a brief period (∼10s) and then a bright light was illuminated. A door to escape the light chamber was then opened to allow free access to a darker chamber after crossing through a small middle tunnel with a floor that contained varying levels of metal spikes. Mice were allowed to complete the task 4 times for a duration of 3 min each. On each trial, the floor of the middle chamber was raised from 0mm to 2mm in 0.5mm increments. Time spent in each chamber was recorded and percent time avoiding the light or mechanical stimulus was determined per mouse and then averaged per group for comparison.

For the choice assay (no light), mice were placed in a three-chamber set up for a 3min for acclimatization. The first chamber was empty, the second chamber contained the varying levels of spikes similar to that described for the MCA, and the third chamber contained bedding from the housing that the mouse resided. For the experiment, the mouse is placed in first chamber for 10sec. A door to escape the first chamber was then opened to allow free access to a bedding chamber, which is provided after crossing through a second chamber that contained varying levels of metal spikes. Mice were allowed to complete the task 4 times for a duration of 3 min each. On each trial, the floor of the middle chamber was raised from 0mm to 2mm in 0.5mm increments. Time spent in each chamber was recorded and the percent time avoiding the first chamber that was devoid of beddings was used for comparison to the control. All behavioral assessments were performed in our groups at 1-2 months, 4-5 months and/or 7-9 months of age.

### *Ex Vivo* Recording Preparation

The *ex vivo* hairy hind paw skin/saphenous nerve/dorsal root ganglion (DRG)/spinal cord (SC) somatosensory system recording preparation was performed as described previously [49]. The intracellular single unit recordings were performed on the L2/L3 DRGs using the quartz microelectrode containing 5% Neurobiotin (Vector Laboratories, Burlingame, CA) in 1M potassium acetate. Electrical stimuli were delivered through a suction electrode from the nerve to identify sensory neuron somata with axons contained in the saphenous nerve. When the cell was found to be electrically driven, the peripheral receptive field (RFs) was localized using a small paintbrush, or hot (∼51°C) or cold (∼1°C) physiological saline if no mechanical RF was found. Once identified, RFs were then probed with an increasing series of Von Frey filaments (0.07g-10g, if mechanically sensitive) for 1-2s to assess mechanical responsiveness.

After mechanical responsiveness was determined, a controlled thermal stimulus was applied using a 3 x 5 mm contact area Peltier element (Yale University Machine Shop, New Haven, CT, USA). Cold stimuli consisted of a variable rate cold ramp beginning at 31°C, dropping to approximately 2°C to 4°C, holding for 4 to 5 s and slowly returning to 31°C. After bath temperature was maintained for approximately 4 to 5 s, a heat ramp was applied which went from 31°C to 52°C in 12 s. This heat stimulus was then held at 52°C for 5 s. The stimulus then ramped back down to 31°C in 12 s. Adequate recovery times (approximately 20–30 s) were employed between stimulations. All elicited responses were recorded digitally for offline analysis of thresholds, firing rates, and mean peak instantaneous frequencies (IFs) to the various stimuli using Spike2 software (Cambridge Electronic Design, Cambridge, UK).

### Immunohistochemistry

DRGs from mice at the indicated time points, were removed and immersion fixed in 3% paraformaldehyde in 0.1M phosphate buffer (PB) for 30 minutes at room temperature. Fixed DRGs were embedded in OCT and incubated at -80°C. DRG sections were cut on a cryostat at 12μm and mounted onto gelatin-coated permafrost slides. Sections were then fixed for ∼15 minutes, blocked and incubated overnight with up to two of the following primary antibodies: transient receptor potential vanilloid type 1 (rabbit anti-TRPV1, Alomone; 1:3000), acid-sensing ion channel 3 (guinea pig anti-ASIC3, Millipore; 1:2000), S100β (rabbit anti- S100β, Abcam 1:1000) or GDNF (rabbit anti-GDNF, Abcam 1:500). Sections were then incubated with appropriate fluorescently conjugated secondary antibodies (Jackson ImmunoResearch anti-guinea pig AlexaFluor647; 1:400 or Jackson ImmunoResearch antirabbit AlexaFluor594; 1:400). Slides were coverslipped in Fluro-Gel (Electron Microscopy Sciences) and stored in the dark at room temperature until imaged. Labeling was characterized and documented using a Nikon confocal microscope with sequential scanning to avoid bleed-through of the different fluorophores. Three images were taken from the three different slides from three different animals along with their respective controls. The final intensity was used to generate the graphs as shown in the result section.

### Electron Microscopy

Mice used for Electron microscopy, were perfusion fixed in a solution combined with 4% paraformaldehyde & 2.5% glutaraldehyde in 0.1-M phosphate buffer at pH. 7.4. Saphenous nerve was dissected out, postfixed in same fixation overnight, then transferred to 0.175 mol/L cacodylate buffer, osmicated, dehydrated, and embedded in Embed 812 (Ladd Research Industries). Semithin sections were cut, and the best block was selected for ultrathin sections. Ultrathin sections were stained in uranyl acetate and lead citrate and viewed on a Hitachi H-7600 microscope. Remak bundles were counted from the photographs and grouped into 1-2, 3-5 & more than 6 Remak bundles and % of Remak bundles was calculated & compared between genotypes.

### Single Cell RNA-Seq

CellChat (http://www.cellchat.org, version 1.6.0) objects were created from 4 Seurat (https://satijalab.org/seurat, version 3.1.2) objects (2- and 7- month old wild-type moue DRG controls, 2-month-old mouse neurofibroma pretumor (DhhCre;Nf1^fl/fl^), 7-month-old mouse neurofibroma tumor (DhhCre;Nf1^fl/fl^)) extracted from the 10x scRNA-seq data [65].

The ‘secreted signaling interactions’ sub-database (for mouse) was chosen to infer the cell state-specific communications. Briefly, CellChat identifies over-expressed ligands or receptors in one cell group and then identify over-expressed ligand-receptor interactions if either ligand or receptor is over-expressed. CellChat infers the biologically significant cell-cell communication by assigning each interaction with a probability value and performing a permutation test. These steps create a complex cell-cell communication networks with assigned the communication probability scores.

After inferring aggregated cell-cell networks, we removed autocrine interactions and focused on cell-cell interactions where Schwann cell (SC) lineage cells and neuron subtypes participate as sources (i.e., ligand-expressing) and targets (i.e. receptor-expressing), respectively. The p-value of 0.05 was chosen to extract significant ligand-receptor (LR) interactions for each sample set. We investigated (1) 2 mo. pretumor (case) vs. 2 mo. control, (2) 7 mo. control (case) vs. 2 mo. control, (3) 7 mo. tumor (case) vs. 2 mo. pretumor, (4) 7 mo. tumor (case) vs. 7 mo. control and extracted unique LR pairs only detected in case sample from each comparison. These unique LR pairs were visualized using circle plots, including Schwann cell lineage and neuron subtypes. The same LR pairs were searched against all cell types and visualized using heatmaps [Fig4]. Neuron subtypes were re-annotated based on [64]. Neuron 1 = low-threshold mechanoreceptors (NF), Neuron 2 = lightly myelinated Aδ nociceptors (PEP2), Neuron 3 = C-type thermo-nociceptors (PEP1), Neuron 4 = C-low threshold mechanoreceptors (TH), Neuron 5 = itch-specific sensory neurons (NP2/3), and Neuron 6 = polymodal nociceptors (NP1).

### Real-Time PCR

DRGs (L2/3) were collected, and RNA isolated using a Qiagen RNeasy® Mini Kit® (QIAGEN, Valencia, CA, USA) according to the manufacturer’s protocol. 1μg samples of purified RNA were reverse transcribed into cDNA with M-MLV Reverse Transcriptase (Promega, Madison, WI, USA). 25ng samples of cDNA were then used in SYBR Green real-time PCR reactions on a Step-One real-time PCR machine (Applied Biosystems). Cycle time (Ct) values for all targets were all normalized to a GAPDH internal control, and fold change was determined as 2^ΔΔCt^ (Applied Biosystems). Values were converted and reported as a percentage change, where 2-fold change = 100% change.

### Statistics

All datasets were analyzed using GraphPad/Prism or SigmaPlot statistical software. Data are represented as mean ± SEM. Significance was defined with *P* ≤ 0.05. For behavioral analyses, 1- or 2-way ANOVA (with or without repeated measures) with Tukey’s or Holm-Sidak (HSD) post hoc tests were performed. For electrophysiological analyses, 1 or 2-way ANOVA (with or without repeated measures) with Tukey’s or HSD post hoc tests were completed. For PCR, one way ANOVA with Tukey’s test were performed. For cell culture and IHC, 1 or 2-way ANOVA with Tukey’s test were completed.

## Conflict of interest

The authors declare no competing financial interests.

## Acknowledgements

This work was supported by grants from the NIH to MPJ (R01 NS105715) and NR (R01 NS28840), a grant from the DAMD to NR (DOD W81XWH-19-1-0816), Young Investigator Awards from the Children’s Tumor Foundation to NGRR (CTF-2022-01-007) and JPC (CTF-2020-01-006), and support from the Cincinnati Children’s Research Foundation the Department of Anesthesia.

---

**Supplementary Figure 1:**
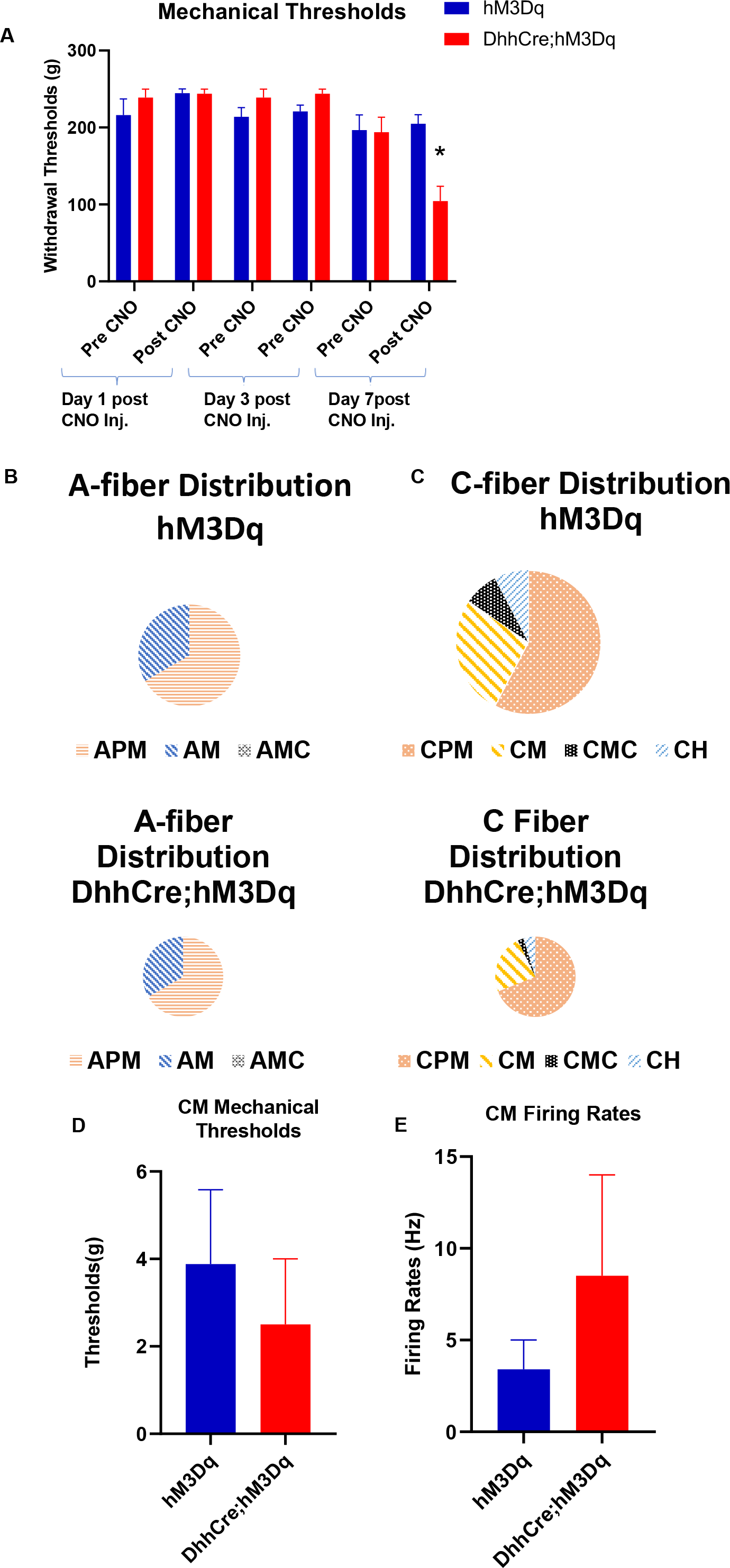
**CNO dose response analysis and additional response properties obtained from DhhCre;hM3Dq mice**. **A:** Injection of CNO for seven days induces peripheral hypersensitivity in hM3Dq mice while no change in the mechanical hypersensitivity is observed after 1 or 3d of CNO (n=5 control, n=5 mutant), *p<0.05, 1-way ANOVA with Tukey’s post hoc). **B:** Distribution of various subtypes of A-fibers found in hM3Dq and DhhCre;hM3Dq mice during *ex vivo* recording; APM: A-polymodal, AM: mechanosensitive only, AMC: A-mechano-cold (n= 2 APM, and 1 AM for control; 2 APM, and 1 AM for mutant). **C** : Distribution of C-fibers in DhhCre;hM3Dq mice compared to controls (hM3Dq); CM: C-mechano, CMC: C-mechano cold, CH: C-heat (n= 7 CM, 2 CMC and 2 CH for control; n= 10 CM, 1 CMC and 2 CH for mutant). **D-F**: The mechanical thresholds and firing rates of CM fibers were not different when compared to the controls (n= 7 hM3Dq, n=9 DhhCre;hM3Dq, p> 0.05 vs hM3Dq, 1-way ANOVA with Tukey’s post hoc; Mean ± SEM).

**Supplementary Figure 2:**
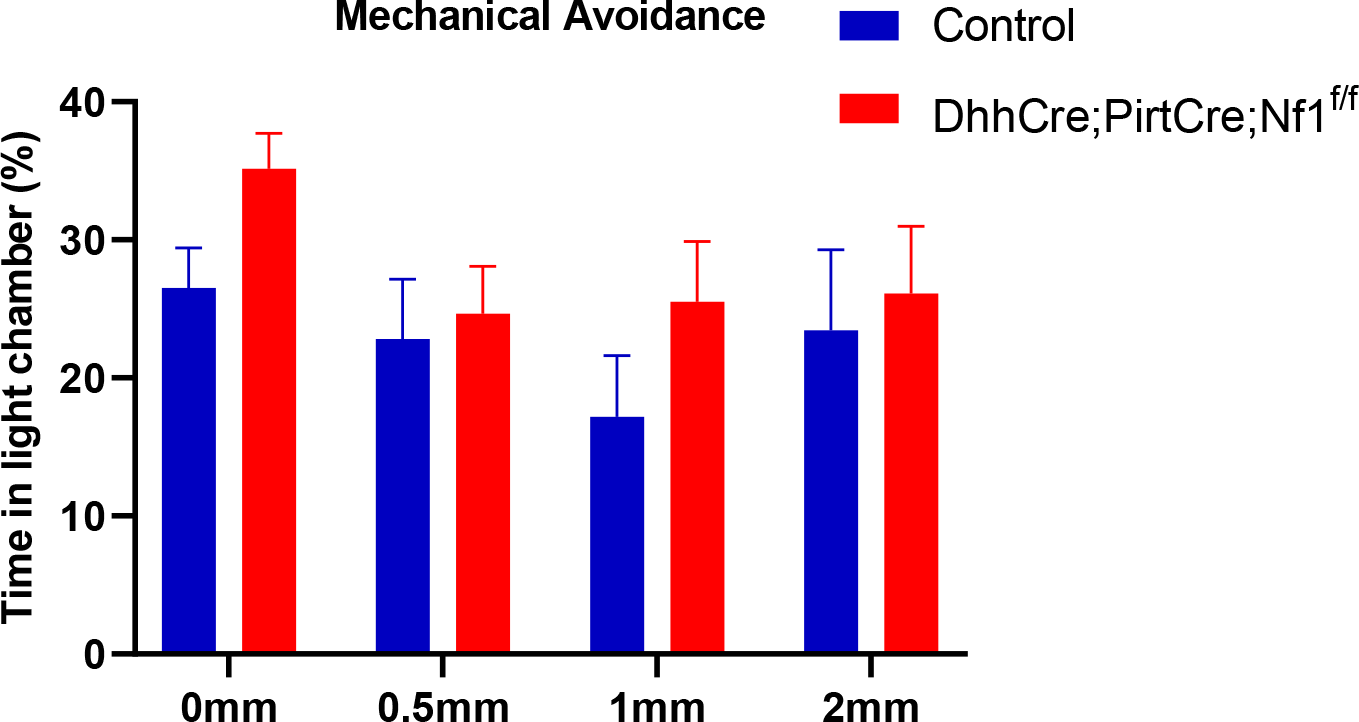
Behavioral analysis of mice with one copy of *Nf1* deleted from both SCs and sensory neurons. No differences in mechanical avoidance are found in mice with one copy deleted from SCs and sensory neurons (DhhCre;PirtCre;Nf1^+/f^) compared to controls (control= 14, mutant= 14, p> 0.05, 2-way ANOVA, Tukey’s post hoc; Mean ± SEM).

**Supplementary Figure 3:**
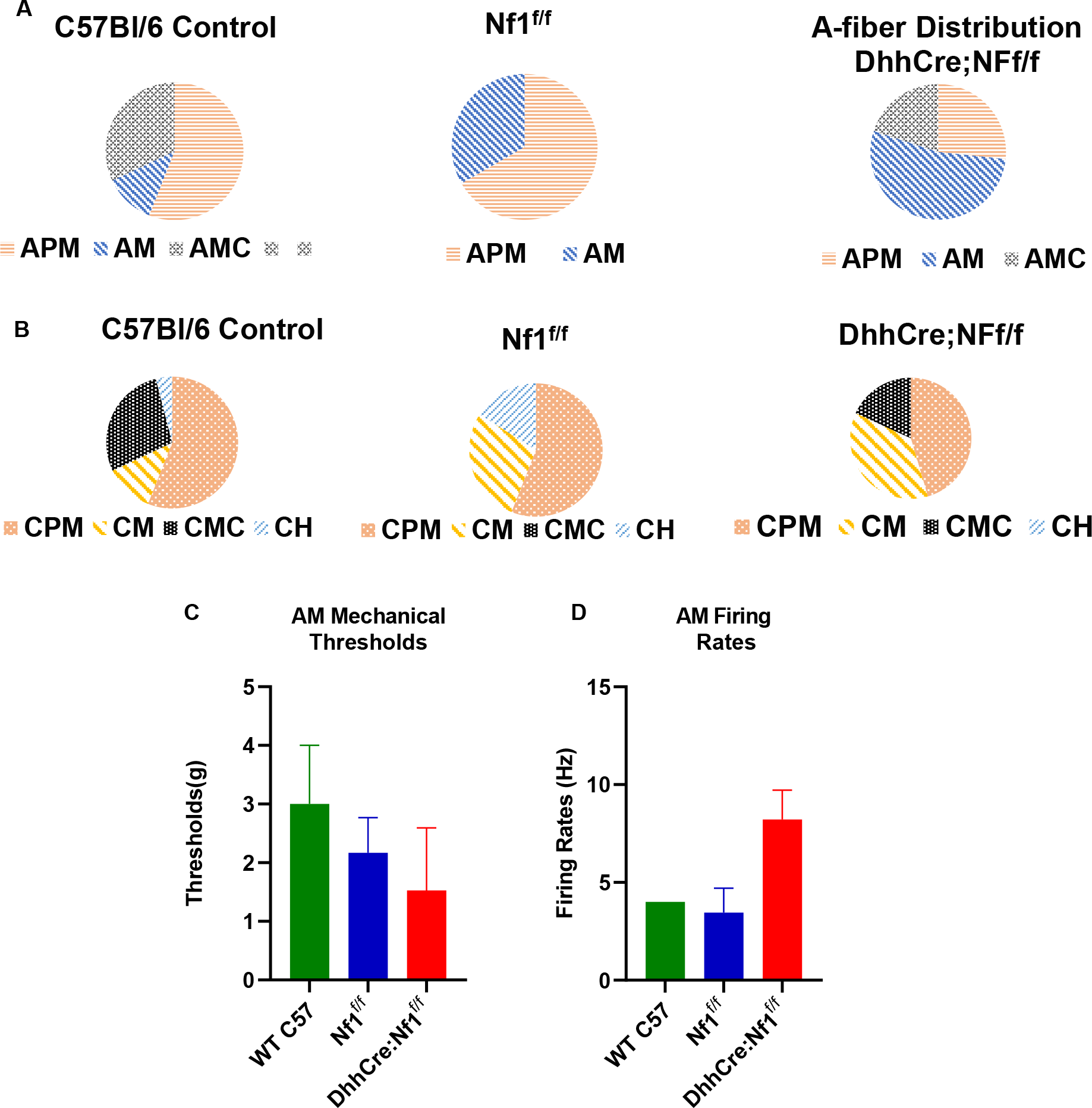
Response properties and distribution of A-fibers and C-fibers in WT C57BI6, Nf1^f/f^ and DhhCre;Nf1^f/f^ mice. A: Distribution of A-fibers in C57Bl6, Nf1^f/f^ and DhhCre;Nf1^f/f^ mice: APM: A-polymodal, AM: mechanosensitive only, AMC: A-mechano-cold (n= 14 APM, 2 M, and 2 AMC for C57BI6, 12 APM, and 6 AM for Nf1^f/f^; n= 4 APM, 8 AM and 3 AMC for DhhCre;Nf1^f/f^). **B:** Distribution of C-fibers in our three groups. CPM: C-polymodal, CM: C-mechano, CMC: C-mechano cold, CH: C-heat (n= 14 CPM, 3 CM, 7 CMC, and 1 CH for C57BI6; n= 14 CPM, 7 CM, and 4 CH for Nf1^f/f^; n= 5 CPM, 4 CM, and 2 CMC DhhCre;Nf1^f/f^). **C-D**: The mechanical thresholds and firing rates of AM fibers were not different in DhhCre;Nf1^f/f^ mice compared to controls (C57Bl6 WT= 2, Nf1^f/f=^6 and DhhCre;Nf1^f/f^=11, p>0.05, 1-way ANOVA, with Tukey’s post hoc; Mean ± SEM).

**Supplementary Figure 4.**
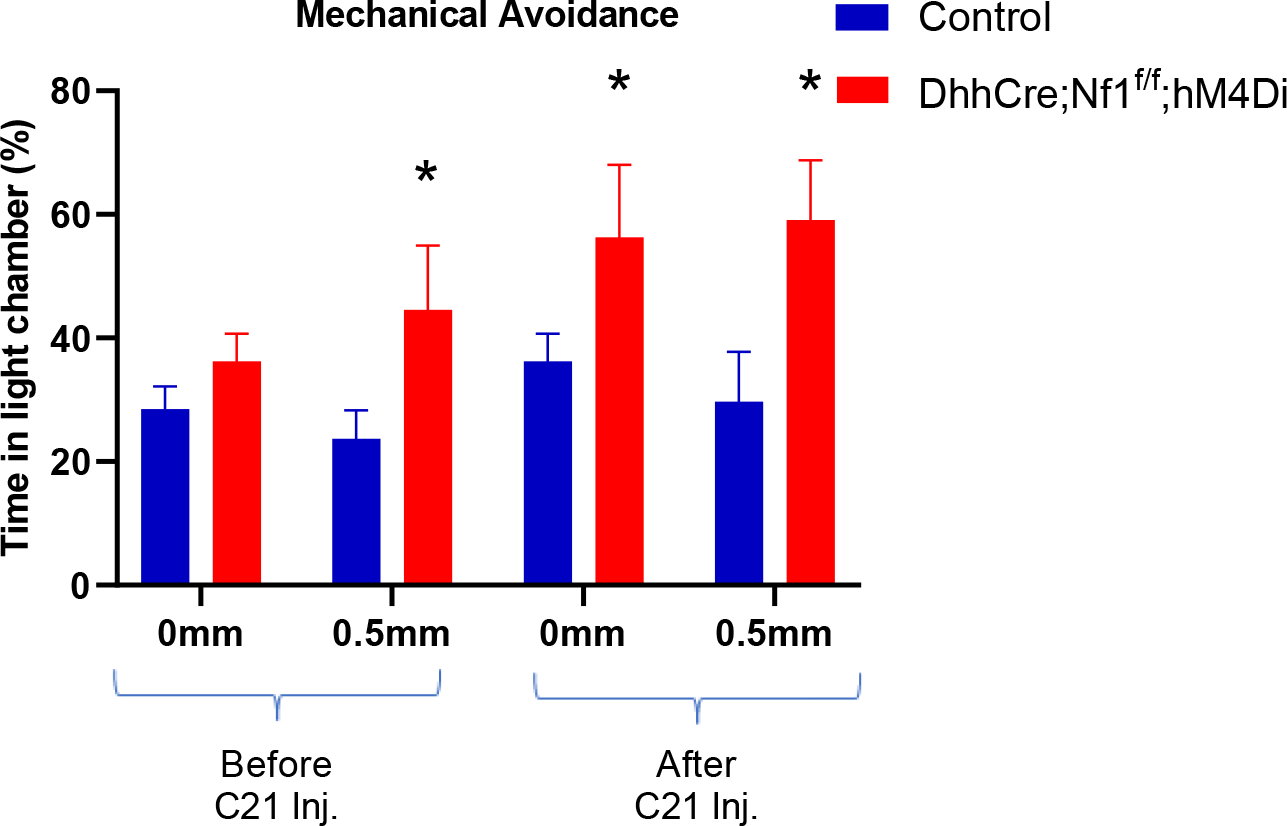
: Mechanical conflict avoidance assay in DhhCre;Nf1^f/f^;hM4Di mice. Prior to C21 treatment, DhhCre;Nf1f/f;hM4Di mice display expected mechanical avoidance, however after 7d of C21 treatment, mice show no avoidance of light and spend approximately equal time in the light and dark chambers regardless of whether the noxious mechanical stimulus is present. (control= 13, mutant= 10, *p<0.05 vs control, 2-way ANOVA with Tukey’s post hoc test; Mean ± SEM).

**Supplementary Figure 5:**
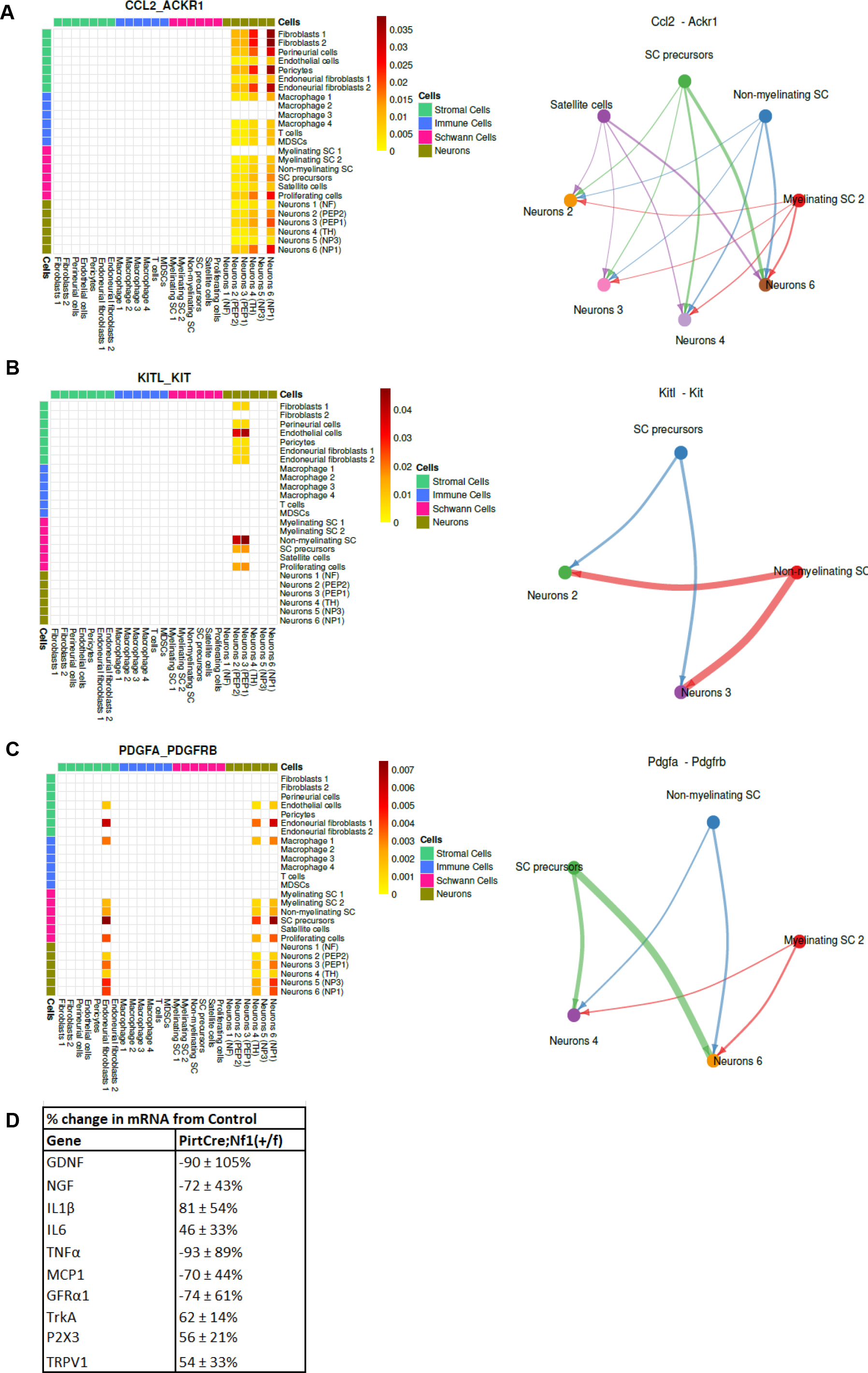
CellChat analysis of additional genes with signaling between glia and sensory neuron subtypes. The heatmaps and signaling diagrams depict CCL2- ACKR1 (A), KITL-KIT (B), and PDGFA- PDGFRB (C) interactions among multiple cell types. These plots suggest that non-myelinating SC, SC precursors, and 4 neuron subtypes display unique interaction partners in 7 mo tumor compared to 2 mo control/pretumor or 7 mo control. D: Realtime PCR from PirtCre;Nf1^+/f^ DRGs shows no differences compared to controls (p>0.05, 1-way ANOVA, Values = % change ± variance).

**Supplementary Figure 6:**
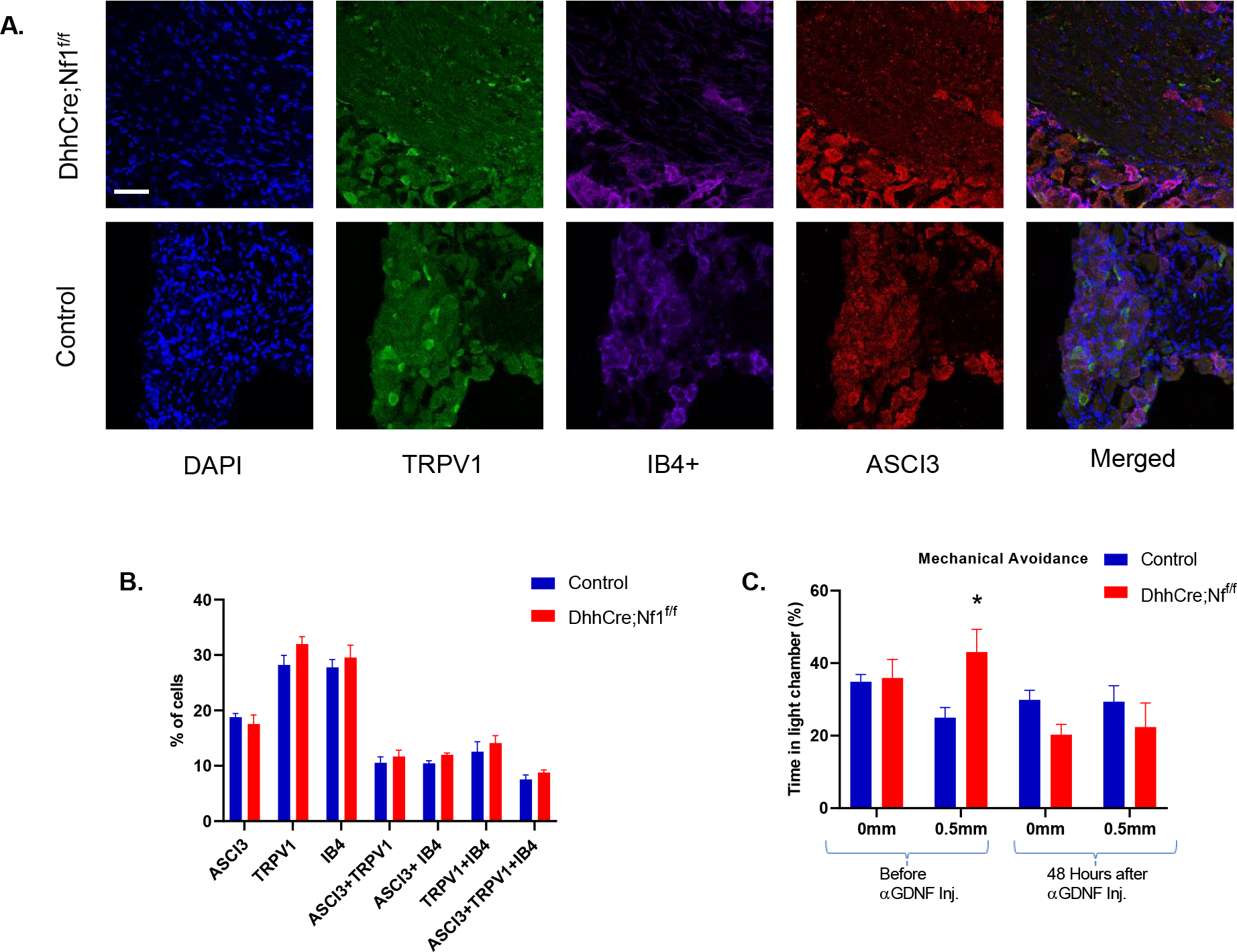
Immunohistochemical analysis of DRGs for various neuronal markers and MCA data from DhhCre;Nf1f/f mice treated with GDNF targeting antibody at 48hrpost injection. A: Examples of DRGs from control or DhhCre;Nf1^f/f^ mice stained for TRPV1, IB4 or ASIC3. B: Quantification of numbers of cells positive for each marker. No differences in numbers of TRPV1, IB4or ASIC3 cells are found in DhhCre;Nf1^f/f^ mice compared to controls (p>0.05, 1-way ANOVA; Mean ± SEM). C: GDNF targeting antibody inhibited mechanical hypersensitivity observed in DhhCre;Nf1f/f mice for at least 48hrs. (n=19 control; n=8 mutant, *p<0.05, 2-way ANOVA, with Tukey’s post hoc; Mean ± SEM).

**Supplementary Table 1:**
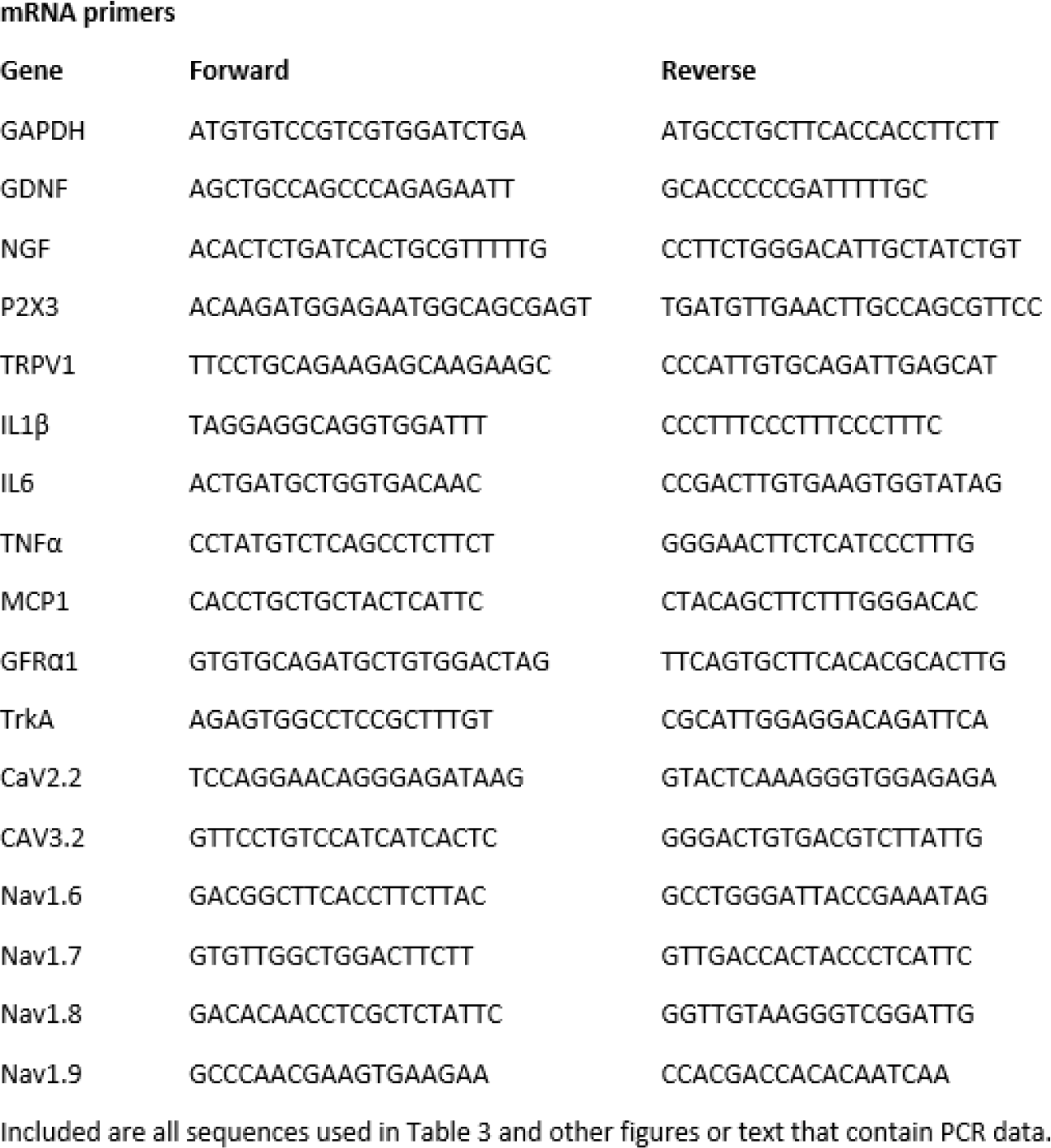
Primer information for gene tested using realtime PCR.

## Notes

### Competing Interest Statement

The authors have declared no competing interest.

## References

1. Adelman, P.C., et al., Single-cell q-PCR derived expression profiles of identified sensory neurons. Mol Pain, 2019. 15: p. 1744806919884496.

2. Baron, R., A. Binder, and G. Wasner, Neuropathic pain: diagnosis, pathophysiological mechanisms, and treatment. Lancet Neurol, 2010. 9(8): p. 807–19.

3. Basbaum, A.I., et al., Cellular and molecular mechanisms of pain. Cell, 2009. 139(2): p. 267–84.

4. Boada, M.D., T.J. Martin, and D.G. Ririe, Nerve injury induced activation of fast-conducting high threshold mechanoreceptors predicts non-reflexive pain related behavior. Neurosci Lett, 2016. 632: p. 44–9.

5. Abdo, H., et al., Specialized cutaneous Schwann cells initiate pain sensation. Science, 2019. 365(6454): p. 695-699.

6. Ji, R.R., T. Berta, and M. Nedergaard, Glia and pain: is chronic pain a gliopathy? Pain, 2013. 154 **Suppl 1**(0 1): p. S10-s28.

7. Wei, Z., et al., Emerging Role of Schwann Cells in Neuropathic Pain: Receptors, Glial Mediators and Myelination. Front Cell Neurosci, 2019. 13: p. 116.

8. Koyanagi, M., et al., Pronociceptive Roles of Schwann Cell-Derived Galectin-3 in Taxane-Induced Peripheral Neuropathy. Cancer Res, 2021. 81(8): p. 2207–2219.

9. Bellampalli, S.S. and R. Khanna, Towards a neurobiological understanding of pain in neurofibromatosis type 1: mechanisms and implications for treatment. Pain, 2019. 160(5): p. 1007–1018.

10. Singh, J.A., S. Noorbaloochi, and K.L. Knutson, Cytokine and neuropeptide levels are associated with pain relief in patients with chronically painful total knee arthroplasty: a pilot study. BMC Musculoskelet Disord, 2017. 18(1): p. 17.

11. Rasmussen, S.A. and J.M. Friedman, NF1 gene and neurofibromatosis 1. Am J Epidemiol, 2000. 151(1): p. 33–40.

12. Legius, E., et al., Revised diagnostic criteria for neurofibromatosis type 1 and Legius syndrome: an international consensus recommendation. Genet Med, 2021. 23(8): p. 1506–1513.

13. Fletcher, J.S., J. Pundavela, and N. Ratner, After Nf1 loss in Schwann cells, inflammation drives neurofibroma formation. Neuro-oncology advances, 2019. 2(Suppl 1): p. i23–i32.

14. White, S., et al., Heat hyperalgesia and mechanical hypersensitivity induced by calcitonin gene-related peptide in a mouse model of neurofibromatosis. PLoS One, 2014. 9(9): p. e106767.

15. Friedman, J.M., Neurofibromatosis 1, in *GeneReviews(®)*, M.P. Adam, et al., Editors. 1993, University of Washington, Seattle Copyright © 1993-2022, University of Washington, Seattle. GeneReviews is a registered trademark of the University of Washington, Seattle. All rights reserved.: Seattle (WA).

16. Prada, C.E., et al., Neurofibroma-associated macrophages play roles in tumor growth and response to pharmacological inhibition. Acta neuropathologica, 2013. 125(1): p. 159–168.

17. Tucker, T., et al., Association between benign and malignant peripheral nerve sheath tumors in NF1. Neurology, 2005. 65(2): p. 205–11.

18. Korf, B.R., Plexiform neurofibromas. Am J Med Genet, 1999. 89(1): p. 31–7.

19. Canavese, F. and J.I. Krajbich, Resection of plexiform neurofibromas in children with neurofibromatosis type 1. J Pediatr Orthop, 2011. 31(3): p. 303–11.

20. Kongkriangkai, A.M., et al., Substantial pain burden in frequency, intensity, interference and chronicity among children and adults with neurofibromatosis Type 1. Am J Med Genet A, 2019. 179(4): p. 602–607.

21. Moutal, A., et al., CRISPR/Cas9 editing of Nf1 gene identifies CRMP2 as a therapeutic target in neurofibromatosis type 1-related pain that is reversed by (S)-Lacosamide. Pain, 2017. 158(12): p. 2301–2319.

22. McCormick, F., Ras signaling and NF1. Curr Opin Genet Dev, 1995. 5(1): p. 51–5.

23. Collins, F.S., et al., Progress towards identifying the neurofibromatosis (NF1) gene. Trends Genet, 1989. 5(7): p. 217–21.

24. Simanshu, D.K., D.V. Nissley, and F. McCormick, RAS Proteins and Their Regulators in Human Disease. Cell, 2017. 170(1): p. 17–33.

25. Friedrich, R.E., et al., Malignant peripheral nerve sheath tumors (MPNST) in neurofibromatosis type 1 (NF1): diagnostic findings on magnetic resonance images and mutation analysis of the NF1 gene. Anticancer Res, 2005. 25(3a): p. 1699–702.

26. Serra, E., et al., Confirmation of a double-hit model for the NF1 gene in benign neurofibromas. Am J Hum Genet, 1997. 61(3): p. 512–9.

27. Kluwe, L., R.E. Friedrich, and V.F. Mautner, Allelic loss of the NF1 gene in NF1-associated plexiform neurofibromas. Cancer Genet Cytogenet, 1999. 113(1): p. 65–9.

28. Riva, M., et al., Recurrent NF1 gene variants and their genotype/phenotype correlations in patients with Neurofibromatosis type I. Genes Chromosomes Cancer, 2022. 61(1): p. 10–21.

29. Corsello, G., et al., Clinical and molecular characterization of 112 single-center patients with Neurofibromatosis type 1. Ital J Pediatr, 2018. 44(1): p. 45.

30. Ratner, N. and S.J. Miller, A RASopathy gene commonly mutated in cancer: the neurofibromatosis type 1 tumour suppressor. Nat Rev Cancer, 2015. 15(5): p. 290–301.

31. Li, H., et al., Immortalization of human normal and NF1 neurofibroma Schwann cells. Lab Invest, 2016. 96(10): p. 1105–15.

32. Jessen, K.R. and R. Mirsky, Schwann Cell Precursors; Multipotent Glial Cells in Embryonic Nerves. Front Mol Neurosci, 2019. 12: p. 69.

33. Pennanen, P., et al., The effect of estradiol, testosterone, and human chorionic gonadotropin on the proliferation of Schwann cells with NF1 (+/-) or NF1 (-/-) genotype derived from human cutaneous neurofibromas. Mol Cell Biochem, 2018. 444(1-2): p. 27–33.

34. O’Brien, D.E., et al., Assessment of pain and itch behavior in a mouse model of neurofibromatosis type 1. J Pain, 2013. 14(6): p. 628–37.

35. Moutal, A., et al., CRMP2-Neurofibromin Interface Drives NF1-related Pain. Neuroscience, 2018. 381: p. 79–90.

36. Cai, S., et al., Targeting T-type/CaV3.2 channels for chronic pain. Transl Res, 2021. 234: p. 20–30.

37. Ferner, R.E., et al., Guidelines for the diagnosis and management of individuals with neurofibromatosis 1. J Med Genet, 2007. 44(2): p. 81–8.

38. Radomska, K.J., et al., Cellular Origin, Tumor Progression, and Pathogenic Mechanisms of Cutaneous Neurofibromas Revealed by Mice with Nf1 Knockout in Boundary Cap Cells. Cancer Discov, 2019. 9(1): p. 130–147.

39. Wu, J., et al., Plexiform and dermal neurofibromas and pigmentation are caused by Nf1 loss in desert hedgehog-expressing cells. Cancer Cell, 2008. 13(2): p. 105–16.

40. Patritti-Cram, J., et al., Purinergic signaling in peripheral nervous system glial cells. Glia, 2021. 69(8): p. 1837–1851.

41. Barnard, E.A., G. Burnstock, and T.E. Webb, G protein-coupled receptors for ATP and other nucleotides: a new receptor family. Trends in Pharmacological Sciences, 1994. 15(3): p. 67–70.

42. Jankowski, M.P., et al., Sensitization of cutaneous nociceptors after nerve transection and regeneration: possible role of target-derived neurotrophic factor signaling. J Neurosci, 2009. 29(6): p. 1636–47.

43. Li, C.-L., et al., Somatosensory neuron types identified by high-coverage single-cell RNA-sequencing and functional heterogeneity. Cell research, 2016. 26(1): p. 83–102.

44. Rice, F.L., et al., The evolution and multi-molecular properties of NF1 cutaneous neurofibromas originating from C-fiber sensory endings and terminal Schwann cells at normal sites of sensory terminations in the skin. PLoS One, 2019. 14(5): p. e0216527.

45. Roth, B.L., DREADDs for Neuroscientists. Neuron, 2016. 89(4): p. 683–694.

46. Gomez, J.L., et al., Chemogenetics revealed: DREADD occupancy and activation via converted clozapine. Science, 2017. 357(6350): p. 503-507.

47. Wu, J., et al., Plexiform and dermal neurofibromas and pigmentation are caused by Nf1 loss in desert hedgehog-expressing cells. Cancer cell, 2008. 13(2): p. 105–116.

48. Jankowski, M.P., et al., Age-dependent sensitization of cutaneous nociceptors during developmental inflammation. Mol Pain, 2014. 10: p. 34.

49. Lu, P., et al., Upregulation of P2Y1 in neonatal nociceptors regulates heat and mechanical sensitization during cutaneous inflammation. Mol Pain, 2017. 13: p. 1744806917730255.

50. Tsuda, M., P2 receptors, microglial cytokines and chemokines, and neuropathic pain. J Neurosci Res, 2017. 95(6): p. 1319–1329.

51. Kiguchi, N., Y. Kobayashi, and S. Kishioka, Chemokines and cytokines in neuroinflammation leading to neuropathic pain. Curr Opin Pharmacol, 2012. 12(1): p. 55–61.

52. Zhang, J.-M. and J. An, Cytokines, inflammation, and pain. International anesthesiology clinics, 2007. 45(2): p. 27–37.

53. Korf, B.R., Neurofibromatosis. Handb Clin Neurol, 2013. 111: p. 333–40.

54. Gutmann, D.H., et al., Neurofibromatosis type 1. Nat Rev Dis Primers, 2017. 3: p. 17004.

55. Santos-Nogueira, E., et al., Randall-Selitto test: a new approach for the detection of neuropathic pain after spinal cord injury. Journal of neurotrauma, 2012. 29(5): p. 898–904.

56. Viisanen, H., et al., Novel RET agonist for the treatment of experimental neuropathies. Mol Pain, 2020. 16: p. 1744806920950866.

57. Zhou, S. and S. Carlton, A novel operant method testing acute cold hypersensitivity in mice using a modification of the Coy Mechanical Conflict-Avoidance System™. The Journal of Pain, 2012. 13(4, Supplement): p. S46.

58. Carlton, S. and S. Zhou, A novel operant method to test acute heat hypersensitivity in mice using a modification of the Coy Operant Mechanical Conflict Avoidance System. The Journal of Pain, 2013. 14(4, Supplement): p. S43.

59. Tran, F.H., et al., Does chronic systemic injection of the DREADD agonists clozapine-N-oxide or Compound 21 change behavior relevant to locomotion, exploration, anxiety, and depression in male non-DREADD-expressing mice? Neurosci Lett, 2020. 739: p. 135432.

60. Armbruster, B.N., et al., Evolving the lock to fit the key to create a family of G protein-coupled receptors potently activated by an inert ligand. Proc Natl Acad Sci U S A, 2007. 104(12): p. 5163–8.

61. Fitzgerald, M. and R. McKelvey, Nerve injury and neuropathic pain - A question of age. Experimental neurology, 2016. 275 **Pt** **2**: p. 296–302.

62. Luo, C., et al., Peripheral Brain Derived Neurotrophic Factor Precursor Regulates Pain as an Inflammatory Mediator. Scientific Reports, 2016. 6(1): p. 27171.

63. Xie, W.R., et al., Robust increase of cutaneous sensitivity, cytokine production and sympathetic sprouting in rats with localized inflammatory irritation of the spinal ganglia. Neuroscience, 2006. 142(3): p. 809–22.

64. Parpaite, T., et al., Patch-seq of mouse DRG neurons reveals candidate genes for specific mechanosensory functions. Cell Rep, 2021. 37(5): p. 109914.

65. Kershner, L.J., et al., Multiple Nf1 Schwann cell populations reprogram the plexiform neurofibroma tumor microenvironment. JCI Insight, 2022. 7(18).

66. Usoskin, D., et al., Unbiased classification of sensory neuron types by large-scale single-cell RNA sequencing. Nat Neurosci, 2015. 18(1): p. 145–53.

67. Jessen, K.R., Glial cells. Int J Biochem Cell Biol, 2004. 36(10): p. 1861–7.

68. Qu, W.R., et al., Interaction between Schwann cells and other cells during repair of peripheral nerve injury. Neural Regen Res, 2021. 16(1): p. 93–98.

69. Cairns, B.E., L. Arendt-Nielsen, and P. Sacerdote, Perspectives in Pain Research 2014: Neuroinflammation and glial cell activation: The cause of transition from acute to chronic pain? Scand J Pain, 2015. 6(1): p. 3–6.

70. Liao, J.Y., et al., Schwann cells and trigeminal neuralgia. Mol Pain, 2020. 16: p. 1744806920963809.

71. Paricio-Montesinos, R., et al., The Sensory Coding of Warm Perception. Neuron, 2020. 106(5): p. 830–841.e3.

72. Lewin, G.R. and L.M. Mendell, Regulation of cutaneous C-fiber heat nociceptors by nerve growth factor in the developing rat. J Neurophysiol, 1994. 71(3): p. 941–9.

73. Boada, D.M., et al., Fast-conducting mechanoreceptors contribute to withdrawal behavior in normal and nerve injured rats. Pain, 2014. 155(12): p. 2646–2655.

74. Tasdemir-Yilmaz, O.E., et al., Diversity of developing peripheral glia revealed by single-cell RNA sequencing. Dev Cell, 2021. 56(17): p. 2516–2535.e8.

75. Alvarez-Leefmans, F.J., et al., Immunolocalization of the Na(+)-K(+)-2Cl(-) cotransporter in peripheral nervous tissue of vertebrates. Neuroscience, 2001. 104(2): p. 569–82.

76. Campana, W.M., Schwann cells: activated peripheral glia and their role in neuropathic pain. Brain Behav Immun, 2007. 21(5): p. 522–7.

77. Zhang, Z.J., B.C. Jiang, and Y.J. Gao, Chemokines in neuron-glial cell interaction and pathogenesis of neuropathic pain. Cell Mol Life Sci, 2017. 74(18): p. 3275–3291.

78. Hingtgen, C.M., S.L. Roy, and D.W. Clapp, Stimulus-evoked release of neuropeptides is enhanced in sensory neurons from mice with a heterozygous mutation of the Nf1 gene. Neuroscience, 2006. 137(2): p. 637–45.

79. Wang, Y., et al., Sensory neurons from Nf1 haploinsufficient mice exhibit increased excitability. J Neurophysiol, 2005. 94(6): p. 3670–6.

80. Coover, R.A., et al., Tonic ATP-mediated growth suppression in peripheral nerve glia requires arrestin-PP2 and is evaded in NF1. Acta Neuropathol Commun, 2018. 6(1): p. 127.

81. Robinson, J.E., et al., Optical dopamine monitoring with dLight1 reveals mesolimbic phenotypes in a mouse model of neurofibromatosis type 1. Elife, 2019. 8.

82. Rizvi, T.A., et al., A novel cytokine pathway suppresses glial cell melanogenesis after injury to adult nerve. J Neurosci, 2002. 22(22): p. 9831–40.

83. Ribeiro, S., et al., Injury signals cooperate with Nf1 loss to relieve the tumor-suppressive environment of adult peripheral nerve. Cell Rep, 2013. 5(1): p. 126–36.

84. Henderson, C.E., et al., GDNF: a potent survival factor for motoneurons present in peripheral nerve and muscle. Science, 1994. 266(5187): p. 1062-4.

85. Van Raamsdonk, C.D. and M. Deo, Links between Schwann cells and melanocytes in development and disease. Pigment Cell Melanoma Res, 2013. 26(5): p. 634–45.

86. Salvo, E., et al., TNFα promotes oral cancer growth, pain, and Schwann cell activation. Sci Rep, 2021. 11(1): p. 1840.

87. Queme, L.F., et al., A dual role for peripheral GDNF signaling in nociception and cardiovascular reflexes in the mouse. Proc Natl Acad Sci U S A, 2020. 117(1): p. 698–707.

88. García-Fernández, P., et al., Systemic inflammatory markers in patients with polyneuropathies. Front Immunol, 2023. 14: p. 1067714.

89. Zhu, Q., et al., Effects of Pulsed Radiofrequency on Nerve Repair and Expressions of GFAP and GDNF in Rats with Neuropathic Pain. Biomed Res Int, 2021. 2021: p. 9916978.

90. Rau, K.K., et al., Mrgprd enhances excitability in specific populations of cutaneous murine polymodal nociceptors. J Neurosci, 2009. 29(26): p. 8612–9.

91. Choi, K., et al., An inflammatory gene signature distinguishes neurofibroma Schwann cells and macrophages from cells in the normal peripheral nervous system. Sci Rep, 2017. 7: p. 43315.

92. Harte, S.E., et al., Mechanical Conflict System: A Novel Operant Method for the Assessment of Nociceptive Behavior. PLoS One, 2016. 11(2): p. e0150164.

